# Mutations in Rv2983 as a novel determinant of resistance to nitroimidazole drugs in *Mycobacterium tuberculosis*

**DOI:** 10.1101/457754

**Authors:** Dalin Rifat, Si-Yang Li, Thomas Ioerger, Jean-Philippe Lanoix, Jin Lee, Ghader Bashiri, James Sacchettini, Eric Nuermberger

## Abstract

Delamanid represents one of two novel antimicrobial classes approved to treat tuberculosis in over 40 years. Pretomanid is another promising nitroimidazole pro-drug in clinical development. Characterization of the full spectrum of mutations conferring resistance to nitroimidazoles and their related phenotypes in *Mycobacterium tuberculosis* will inform development of suitable genotypic and phenotypic drug susceptibility tests. Here, we used a range of pretomanid doses to select pretomanid-resistant mutants in two pathologically distinct murine TB models. The frequency of spontaneous pretomanid resistance mutations was approximately 10^−5^ CFU. Pretomanid demonstrated dose-dependent bactericidal activity and selective amplification of resistant mutants. Whole genome sequencing of 161 resistant isolates from 47 mice revealed 99 unique mutations, 90% of which were found in 1 of 5 genes previously associated with nitroimidazole activation and resistance. The remaining 10% harbored isolated mutations in *Rv2983*. Complementing an *Rv2983* mutant with a wild-type copy of *Rv2983* restored wild-type susceptibility to pretomanid and delamanid, confirming that loss of *Rv2983* function causes nitroimidazole resistance. By quantifying F_420_ and its precursor Fo in *Mycobacterium smegmatis* overexpressing *Rv2983* and an *M. tuberculosis Rv2983* mutant, we provide evidence that Rv2983 is necessary for F_420_ biosynthesis and nitroimidazole activation, perhaps as the guanylyltransferase CofC. F_420_H_2_-deficient mutants displayed hypersusceptibility to malachite green (MG), a selective decontaminant present in solid media used to isolate and propagate mycobacteria from clinical samples. The wide diversity of mutations causing high-level pretomanid resistance and MG hypersusceptibility of most mutants poses significant challenges to clinical detection of nitroimidazole resistance using either genotypic or phenotypic methods.

**Significance:** Nitroimidazole pro-drugs represent a promising new class of anti-tuberculosis drugs. Reliable methods to assure nitroimidazole susceptibility are critical to assure their optimal use. Yet, the spectrum of nitroimidazole resistance mutations remains incompletely characterized. Using 161 pretomanid-resistant *Mycobacterium tuberculosis* isolates selected in pretomanid-treated mice, we discovered a novel resistance determinant, *Rv2983*, required for cofactor F_420_ biosynthesis and characterized the remarkable diversity of mutations in this and 5 other genes involved in nitroimidazole activation. We show that F_420_H_2_–deficient nitroimidazole-resistant mutants are hypersusceptible to the selective decontaminant malachite green used in solid media to isolate mycobacteria and may evade detection on such media. These results have important implications for development and clinical use of genotypic and phenotypic methods for nitroimidazole susceptibility testing.

## Introduction

Despite decades of efforts to end the global tuberculosis (TB) epidemic, *Mycobacterium tuberculosis* is the leading killer among infectious agents plaguing mankind (*1*). The emergence and spread of multidrug-resistant (MDR) and extensively drug-resistant (XDR) *M. tuberculosis* makes the eradication effort much more difficult because treatment requires administration of more toxic and less effective second‐ and third-line drugs for up to 2 years (*1, 2*). Delamanid and pretomanid are promising new bicyclic 4-nitroimidazole drugs that have shown potential in pre-clinical and clinical studies to shorten and simplify the treatment of TB, including drug-resistant forms (*3-9*). Delamanid received conditional approval by the European Medicines Agency (EMA) to treat MDR-TB in 2014 (*10*) and pretomanid is currently being evaluated in Phase 2/3 clinical trials (ClinicalTrials.gov Identifiers: NCT03338621, NCT02589782, NCT02333799, NCT03086486). Particularly notable is a novel regimen comprised of bedaquiline, pretomanid and linezolid that may represent a highly efficacious oral, short-course regimen for treatment of MDR/XDR-TB (*4*)(ClinicalTrials.gov Identifiers: NCT02333799, NCT03086486).

Two non-exclusive mechanisms of action have been described for these bicyclic 4-nitroimidazole drugs: inhibition of cell wall biosynthesis through inhibition of mycolic acid synthesis and respiratory poisoning through release of nitric oxide during bacterial drug metabolism (*11, 12*). Pretomanid and delamanid are prodrugs that require bioreductive activation of an aromatic nitro group by the 8-hydroxy-5-deazaflavin (coenzyme F_420_)-dependent nitroreductase Ddn in order to exert bactericidal activity (*13*). The reaction involves the transfer of two electrons from the reduced form of F_420_ (F_420_H_2_) produced by an F_420_-dependent glucose-6-phosphate dehydrogenase (Fgd1) (*12, 14*). Therefore, F_420_ biosynthesis and reduction are essential for the activation of delamanid, pretomanid and other nitroimidazole prodrugs. Three genes are identified as essential for F_420_ biosynthesis in *M. tuberculosis* complex (*15, 16*). *fbiC* encodes a 7,8-didemethyl-8-hydroxy-5-deazariboflavin (Fo) synthase that catalyzes the condensation of 5-amino-6-ribitylamino-2,4 (1*H*, 3*H*)-pyrimidinedione and tyrosine to form the F_420_ precursor Fo (*17, 18*). *fbiA* encodes a transferase that is believed to catalyze the transfer of a phospholactyl moiety to Fo to generate F_420_–0, while *fbiB* encodes a F_420_–0:γ-L-glutamyl ligase that catalyzes the sequential addition of a variable number of glutamate residues to F_420_–0 to yield coenzyme F_420_-5 or -6 in mycobacteria (*12*). In the methanogen *Methanocaldococcus jannaschii*, a guanylyltransferase termed CofC is believed to generate an intermediate (L-lactyl-2-diphospho-5’-guanosine,–LPPG) in the F_420_ biosynthesis pathway (*19*). A homologous enzyme, MSMEG_2392, is shown to be necessary for F_420_ synthesis in *Mycobacterium smegmatis* through transposon mutagenesis studies (*20*). An ortholog, *Rv2983*, is present in *M. tuberculosis*. However, the role of MSMEG-2392 and Rv2983 in F_420_ biosynthesis has remained unexplored.

Loss-of-function mutations in *ddn*, *fgd1* and *fbiA-C* causing delamanid and pretomanid resistance are readily selected *in vitro* in *M. tuberculosis* complex (*16, 18, 21-23*). However, the genetic spectrum of mutations emerging during *in vivo* selection has not been characterized. In order to study bacterial genetic, host and pharmacological factors associated with emergence of nitroimidazole resistance *in vivo*, we selected pretomanid-resistant mutants using a wide range of pretomanid doses in two mouse models of TB and characterized them by whole genome sequencing (WGS). Because the lungs of TB patients feature a heterogeneous array of lesion types associated with diversified immune responses and drug penetration (*24, 25*), we used both C3HeB/FeJ mice, which develop caseating lung lesions in response to *M. tuberculosis* infection, and BALB/c mice, which do not, to investigate the impact of these caseating lesions and their associated micro-environments on mutant selection. In the present study, we found that pretomanid-resistant mutants were readily selected by monotherapy in both mouse strains. While the majority of resistant isolates harbored isolated mutations in genes previously associated with nitroimidazole resistance, all the resistant isolates lacking such mutations had mutations in *Rv2983*. We went on to confirm that loss-of-function mutations in *Rv2983* cause high-level pretomanid and delamanid resistance through disruption of F_420_ biosynthesis, supporting the hypothesis that *Rv2983* plays a role similar to *cofC* in *M. tuberculosis*. Furthermore, F_420_H_2_-deficient nitroimidazole-resistant *M. tuberculosis* mutants, including *Rv2983* mutants, were found to be hypersensitive to malachite green (MG), an organic compound used as a selective decontaminant in solid media for culturing *M. tuberculosis*, which may have important implications for their detection in clinical samples.

## Materials and Methods

### Bacterial strains, media, antimicrobials and reagents

Wild type *M. tuberculosis* H37Rv (ATCC 27294) was mouse-passaged, frozen in aliquots and used in all the experiments. The wild type *M. smegmatis* strain mc^2^155 was obtained from the stock in the lab. Unless stated otherwise, Middlebrook 7H9 medium (Difco, BD) supplemented with 10% oleic acid-albumin-dextrose-catalase (OADC) complex (BD), 0.5% glycerol and 0.05% Tween 80 (Sigma-Aldrich) (7H9 broth) was used for cultivation. Middlebrook 7H10 agar and selective 7H11 agar (Difco, BD), prepared from powder and containing 10% OADC and 0.5% glycerol, were used for comparison of strain recovery on commercially available agar plates. Lowenstein Jensen (LJ) slants were purchased from BD. Pretomanid and delamanid were kindly provided by the Global Alliance for TB Drug Development (New York, NY).

### Mouse infection models and pretomanid treatment

All animal procedures were approved by the Animal Care and Use Committee of Johns Hopkins University. Aerosol infections were performed using the Inhalation Exposure System (Glas-col Inc., Terre Haute, IN), as previously described (*26*). Briefly, 6-week-old female BALB/c mice (Charles River, Wilmington, MA) and C3HeB/FeJ mice (Jackson Laboratories Bar Harbor, ME) were infected with a log phase culture of *M. tuberculosis* that was grown in 7H9 broth to O.D.600_nm_= 1.0 and then diluted in the same medium prior to infection to deliver 50-100 CFU to the lungs. Pretomanid was formulated for oral administration as previously described (*27*). Beginning 8 weeks after aerosol infection, mice were randomly allocated into groups and treated once daily (5 days per week) for up to 8 weeks with pretomanid at doses of 10, 30, 100, 300 and 1000 mg/kg. Untreated mice were sacrificed on the day after aerosol infection and on the day of treatment initiation to determine the number of CFU implanted in the lungs and pretreatment CFU counts, respectively. Additional mice were sacrificed after 3 and 8 weeks of treatment to evaluate the treatment response. Serial 10-fold dilutions of lung homogenates were plated on 7H11 agar. Week 8 samples including those from untreated mice were also plated in parallel on 7H11 plates containing 0.25, 1 and 10 μg/ml of pretomanid to quantify the resistant CFU. Plates were incubated at 37°C for 28 days before final CFU counts were determined.

### Whole genome sequencing

For each mouse lung that yielded growth on pretomanid-containing plates, individual colonies and, for a subset of mice, pools of up to 15 colonies, were randomly selected from pretomanid-containing plates and sub-cultured in 7H9 broth prior to extraction of genomic DNA using the cetyltrimethylammonium bromide (CTAB) protocol (*28*) and vortexing (Genegate, Inc.). 2-3 μg of genomic DNA was sheared by a nebulizer to generate DNA fragments. The DNA library was prepared using a genomic DNA sample preparation kit (Illumina, Inc.), in which adapter-ligated DNA fragments were 250-350 bp in length, and carried out on an Illumina Genome Analyzer II (Illumina, Inc). The sequencer was operated in paired-end mode to collect pairs of reads of 51-bp from opposite ends of each fragment. Image analysis and base-calling were done by using the Illumina GA Pipeline software (v0.3). The reads that were generated for each strain were aligned to the reference genome of *M. tuberculosis* H37Rv (*29*). Based on alignment to the corresponding region in the reference genome, single nucleotide polymorphism (SNP), insertion and deletion were identified on the genome of resistant strains by using a contig-building algorithm to construct a local ~200 bp sequence spanning the site of mutagenesis (*30*). Distribution of mutation type and mutation frequency in genes involved in nitroimidazole resistance was calculated by counting the total number of unique mutations isolated from each mouse in the same treatment group.

### Complementation of an *Rv2983* mutation

A 1,044-bp DNA fragment containing the open reading frame (ORF) of the wild type *Rv2983* gene, including 340 bp of 5’-flanking sequence and 59 bp of 3’-flanking sequence, was PCR-amplified from *M. tuberculosis* H37Rv genomic DNA using primers Rv2983-1F and *Rv2983-1R* (Table S1). The *Rv2983* PCR product was ligated into XbaI-digested E. coli-mycobacterium shuttle vector pMH94 (*31*) using NE builder HiFi DNA assembly kit (NE Biolabs) to generate the recombinant pMH94-Rv2983 vector. Similarly, a 388-bp DNA fragment containing the *hsp60* promoter and a 645-bp DNA fragment of *Rv2983* open reading frame were amplified from *M. tuberculosis* H37Rv genomic DNA using primer sets *hsp60*-F and *hsp60-R* and *Rv2983-2F* and *Rv2983-2R*, respectively (Table S1), and ligated into Xbal-digested E. coli-mycobacterium shuttle vector pMH94 to yield *pMH94-hsp60-Rv2983*. A small amount of ligation reaction was transferred into *E. coli* competent cells, followed by DNA sequencing of the inserts in the corresponding recombinants. The recombinants *pMH94-Rv2983* and *pMH94-hsp60-Rv2983* were electroporated into competent cells of *Rv2983* mutant strain BA_101 (B101), harboring an A198P substitution, to enable selection of complemented candidates B101pRv2983 and *B101phsp60-Rv2983* on 7H10 agar containing 25 μg/ml of kanamycin. To confirm the complementation genetically, Southern blotting was performed using a digoxigenin (DIG) DNA labeling and detection kit according to the manufacturer’s protocol (Sigma). Briefly, a 448-bp *Rv2983* probe was generated by addition of DIG-dUTP (Sigma) to PCR reactions containing primer pairs *Rv2983-3F* and *Rv2983-3R* (Table S1). Acc65I-digested (NE biolabs) genomic DNA of the wild type, the B101 mutant and the B101pRv2983 and *B101phsp60*-*Rv2983* complemented strains was separated on agarose gel and transferred onto positively-charged nylon-membrane (GE). After pre-hybridization, the membrane was hybridized with the DIG-labeled *Rv2983* probe at 68°C overnight, followed by addition of anti-DIG alkaline phosphatase conjugate. After stringent washes, the membrane was incubated with the chemiluminescence substrate disodium 3-(4-methoxyspiro {1,2-dioxetane-3,2(5′-chloro)tricycloecan}-4-yl)phenyl phosphate (CSPD) and exposed on X-ray film in a dark room prior to development using a developer (AFP imaging)(*32*).

### MIC determination

Log-phase cultures were diluted to achieve a bacterial density of approximately 10^5^ CFU/ml in conical tubes containing 7H9 broth without Tween 80. Serial 10-fold dilutions were plated on 7H11 agar containing stepwise 2-fold increasing pretomanid concentrations ranging from 0.015 to 64 μg/ml or delamanid concentrations from 0.001 to 1.024 μg/ml. Drugs were initially dissolved in dimethylsulfoxide (DMSO) (Sigma) prior to further dilution in 7H9 broth or 7H11 agar. Cultures were incubated at 37°C for 14 days or 28 days after plating. MIC was defined as the lowest drug concentration that inhibited visible *M. tuberculosis* growth in conical tubes or that inhibited 99% of CFU growth on pretomanid-containing plates (*33, 34*). The experiments were repeated twice.

### Construction of recombinants overexpressing *Rv2983*, with or without *fbiC*, in *M. smegmatis*

A 645-bp DNA fragment containing the *Rv2983* ORF was PCR-amplified from *M. tuberculosis* H37Rv genomic DNA using primers Rv2983-4F and *Rv2983-4R* (Table S1). The amplified PCR product was ligated into the NdeI‐ and PacI-digested E. coli-mycobacterium shuttle vector pYUBDuet (*35*) using NE builder HiFi DNA assembly kit (NE Biolabs) and then transferred into Turbo-competent *E. coli* cells (NE Biolabs) prior to plating on LB agar plates containing 100 μg/ml of hygromycin B for selection of recombinants. The *Rv2983* PCR product was also similarly ligated into the same NdeI‐ and PacI-digested pYUBDuet vector harboring *fbiC* (termed *pfbiC*) (*35*) to overexpress both *Rv2983* and *fbiC*. After confirmation by restriction digestion and DNA sequencing, the constructs were electroporated into competent *M. smegmatis* cells prior to selecting recombinants on 7H10 agar plates containing 100 μg/ml of hygromycin B. PCR amplification was used to confirm the inserts on the *M. smegmatis* genome. pYUBDuet and pYUBDuet harboring *fbiA*, *fbiB* and *fbiC* (termed *pfbiABC*) (*35*) were also transferred into competent *M. smegmatis* cells to serve as controls.

### Measurement of Fo and F_420_

Extraction of Fo and F_420_ was performed in *M. smegmatis* and *M. tuberculosis* strains according to a previous study (*35*), with minor modifications. Briefly, *M. smegmatis* strains harboring different constructs and pYUBDuet were grown in 7H9 broth in a shaker to mid-log phase (O.D.600_nm_ = 0.7-1.0), followed by induction using 1mM isopropyl β-D-1-thiogalactopyranoside (IPTG) for 6 and 26 hours. After centrifugation for 15 min at 16000 x g, the supernatants were removed for detection of Fo, which is principally found in culture supernatant whereas F_420_ with 5 or 6 glutamate residues is largely retained inside cells (*15, 35, 36*). The cell pellets were washed with 25mM sodium phosphate buffer (pH 7.0) and re-suspended at 100 mg/mL in the same buffer, then autoclaved at 121°C for 15 min. After centrifugation at 16000 x g for 15 min at 4°C, the cell extracts were harvested for detection of F_420_ (*35*). Fluorescence of the supernatant and cell extracts was measured using an excitation wavelength of 410 nm and an emission wavelength of 465 nm. Fluorescent signals of Fo were normalized using the O.D. at 600_nm_. The small portion of Fo (1-7%) retained inside cells was ignored when quantifying F_420_ in cell extracts (*37*). Relative fluorescent signals were calculated in *M. smegmatis* harboring each of recombinants relative to pYUBDuet alone. Similarly, cell extracts and supernatant were extracted from *M. tuberculosis* strains grown in 7H9 broth for 6 days at initial O.D.600_nm_ of 0.1. Relative fluorescent signals of F_420_ and Fo were calculated using cell extracts and supernatant relative to 25 mM phosphate buffer and 7H9 broth, respectively. *M. smegmatis* harboring pYUBDuet-*fbiABC* was used as a positive signal control for Fo and F_420_ due to their commercial unavailability (*35*). The experiment was repeated twice.

### Quantification of gene expression in *M. tuberculosis*

6-day-old *M. tuberculosis* strains grown in 7H9 broth as described above were sub-cultured in fresh 7H9 at O.D.600_nm_ = 0.05 followed by incubation at 37°C in a shaker for 2 and 4 days. Bacterial pellets were collected by centrifugation at 3500 rpm at 4°C for purification of total RNA using Trizol (Thermo Fisher Scientific) according to the manufacturer’s protocol followed by removal of DNA contamination with Turbo DNAse (Ambion). Following cDNA synthesis with random hexamers and oligo(dT)20 primer and superscript III reverse transcriptase (Invitrogen), quantitative PCR was performed to measure gene expression of *M. tuberculosis* using SYBR Green PCR master mix (Thermo scientific) and StepOne™ system (Applied biosystems) with primer sets listed in Table S1. The cycle threshold value (C_T_) measured for each gene was normalized to that of the housekeeping gene *sigA* (ΔC_T_) amplified by the primers *sigA-F* and *sigA-R* (Table S1). ΔΔC_T_ was calculated in each of pretomanid-resistant strains relative to the wild-type H37Rv prior to calculation of the fold-change in gene expression (2^-ΔΔC_T_) (*38*). All samples were prepared in duplicate. PCR was performed from an equal amount of cDNA samples synthesized with oligo(dT)20 with primers *fbiC-5-7_F* and *fbiC-5-7_R* (Table S1) using the Q_5_ High-fidelity PCR kit (New England Biolabs) and C1000 Thermal cycler (Biorad). The PCR product was examined by electrophoresis on a 1% agarose gel.

### Malachite green susceptibility testing

7H9 media supplemented with 10% OADC, 0.5% glycerol, 1.5% Bacto™ Agar (BD) and malachite green (MG) oxalate (Alfa Aesar) was used to prepare solid 7H9 media with differing MG concentrations. *M. tuberculosis* strains were grown to mid-log phase and diluted to OD600_nm_ = 0.1 in 7H9 broth before serial 10-fold dilutions were plated in 100 or 500 μl aliquots on 7H9 agar containing MG concentrations of 0, 0.1, 0.3, 1, 3, 10, 30, 100, 300, 1000 μg/ml or 0, 3, 6, 12 μg/ml. CFU were counted after 28, 35 and 49 days of incubation. The same cultures were also plated on 7H10 and 7H11 agar plates and LJ slants. Serially diluted cultures were inoculated onto LJ slants using calibrated disposable inoculating loops (10 μl per loop, BD) as one loop per LJ slant. Plates were incubated at 37°C for 21, 28 and 35 days for CFU counts. Colony size was observed weekly until day 35, beginning 21 days after plating. The experiment was repeated two times under similar conditions.

### Statistical analysis

Log_10_-transformed CFU counts, fold-change values of gene expression and absorbance (A_410_) values of fluorescent signals were used to calculate means and standard deviations for each data set. Differences between means were compared by the Student’s *t* test in Microsoft Excel. Differences in mutation frequencies between two mouse models were evaluated by Fisher’s exact test in GraphPad Prism 6. A *p*-value of < 0.05 was considered statistically significant.

## Results

### Spontaneous pretomanid-resistant mutants exist at a relatively high frequency in infected mice and are selectively amplified by treatment with active doses of pretomanid

To study the dose-response of pretomanid and explore the genetic spectrum of nitroimidazole resistance selected *in vivo*, we established chronic *M. tuberculosis* infections in BALB/c and C3HeB/FeJ mice and then treated with a range of pretomanid doses for up to 8 weeks. Despite lower CFU counts on the day after infection (W-8) in C3HeB/FeJ mice (1.67 log_10_ CFU per lung) compared to BALB/c (2.26 log_10_) (*p* <0.001), higher CFU counts were observed in C3HeB/FeJ mice 8 weeks later on the day treatment started (D0) and after 3 weeks of treatment in almost all groups (*p* <0.001 - 0.05) (Fig. 1A). Three C3HeB/FeJ mice treated with 1000 mg/kg required euthanasia during the second week of treatment, prompting a dose reduction from 1000 mg/kg to 600 mg/kg in both strains. Nevertheless, a clear pretomanid dose-response relationship was observed in both mouse strains after 3 weeks of treatment (Fig. 1A). The three remaining C3HeB/FeJ mice treated with 600 mg/kg beyond the week 3 time point were euthanized after 5 weeks of treatment due to toxicity. One had no detectable CFU and two had ≤ 2.0 logi0CFU of pretomanid-resistant *M. tuberculosis*. After 8 weeks of treatment, total CFU counts fell in a dose-dependent manner in BALB/c mice before a plateau was reached at doses ≥ 300 mg/kg, where resistant CFU were higher and replaced the susceptible CFU (*p* < 0.05) (Fig. 1B). Spontaneous pretomanid-resistant CFU comprised approximately 10^−5^of the total CFU in the absence of drug pressure in untreated BALB/c mice and the proportion of the total CFU that was comprised of pretomanid-resistant CFU increased with dose up to the 300 mg/kg dose group. Dose-dependent bactericidal activity was also observed in C3HeB/FeJ mice (Fig. 1C). However, selective amplification of pretomanid-resistant mutants was more extensive and occurred at lower doses than in BALB/c mice (Fig. 1B and 1C). We were not able to measure the spontaneous frequency of resistant mutants in untreated C3HeB/FeJ mice because they succumbed to infection prior to week 8. Pretomanid-resistant CFU replaced susceptible CFU in C3HeB/FeJ mice receiving doses as low as 30 mg/kg and pretomanid-resistant CFU counts were roughly 10 times higher in C3HeB/FeJ mice compared to BALB/c mice (Fig. 1B and C), which indicates greater potential for selective amplification of pretomanid resistance with monotherapy in this strain. Most resistant isolates grew on plates containing 10 μg/ml of pretomanid, but some had fewer CFU on plates containing 10 μg/ml than on those containing 1 μg/ml of pretomanid.

**Fig. 1.**
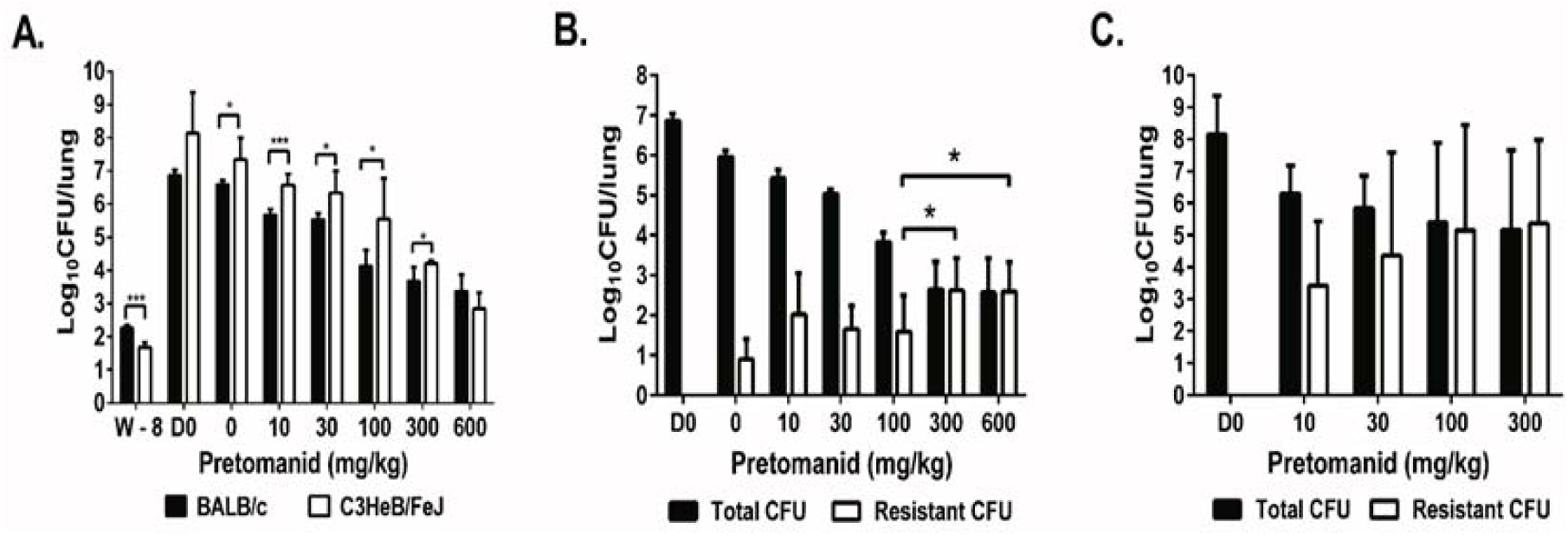
Selective amplification of spontaneous pretomanid-resistant mutants during pretomanid monotherapy in mice is dose-dependent and is more pronounced in C3HeB/FeJ mice. After aerosol infection with *M. tuberculosis* H37Rv, BALB/c and C3HeB/FeJ mice were treated with a range of doses of pretomanid for 8 weeks and sacrificed at different time points before and after treatment for lung CFU counts. A. Mean (± S.D.) total lung CFU counts on the day after infection (W-8), on the day of treatment initiation (D0), and after 3 weeks of treatment with the indicated pretomanid dose (in mg/kg body weight). Dose-dependent bactericidal activity was observed in both strains; B. Mean (± S.D.) total and PMD-resistant lung CFU counts in BALB/c mice on day 0 and after 8 weeks of treatment with the indicated pretomanid dose. Dose-dependent bactericidal activity and selection of PMD-resistant bacteria was observed, with the resistant population overtaking the susceptible population at doses ≥ 300 mg/kg; C. Mean (± S.D.) total and PMD-resistant lung CFU counts in C3HeB/FeJ mice on day 0 and after 8 weeks of treatment with the indicated pretomanid dose. Dose-dependent bactericidal activity and selection of PMD-resistant bacteria was observed, with the resistant population overtaking the susceptible population at doses ≥ 30 mg/kg. * *p*< 0.05, \*\*\**p* < 0.001

### Whole genome sequencing of pretomanid-resistant mutants revealed diverse mutations in *Rv2983* or in one of five other genes required for pretomanid activation

To characterize mutations associated with pretomanid resistance *in vivo*, we performed WGS on 136 individual pretomanid-resistant colonies and 25 colony pools picked from 47 individual mice harboring pretomanid-resistant CFU after 8 weeks of treatment (Table S2 and S3). Each individual isolate had an isolated mutation in *Rv2983* or one of the 5 genes previously shown to be required for pretomanid activation. Overall, 99 unique mutations in these 6 genes were identified from individual and pooled isolates (Table 1 and 2). Except for mutations K9N (*fgd1*), R322L (*fbiC*) and Q120P (*fbiA*), which were shared by two mice each, no two mice harbored the same mutation, which emphasizes the large target size for resistance-conferring mutations. In both BALB/c and C3HeB/FeJ mice, more than half of the resistant isolates were *fbiC* mutants (54 and 56%, respectively) (Table 3). For the other five genes, the rank order by mutation frequency was *Rv2983* (15%) > *fbiA* (13%) > *ddn* (9%) > *fbiB* (6%) > *fgd1* (4%) in BALB/c mice and *fbiA* (18%) > *ddn* (16%) > *fgd1* or *Rv2983* (4%) > *fbiB* (2%) in C3HeB/FeJ mice. No significant differences in mutation frequencies between BALB/c and C3HeB/FeJ mice were observed, although a trend towards more *Rv2983* mutations in BALB/c mice (8/54, 15% of all mutations) compared to C3HeB/FeJ mice (2/45, 4%) was detected. The mutations identified in *Rv2983* included 8 point mutations resulting in the following amino acid substitutions: R25S, R25G, A68E, A132V, G147C, C152R, Q114R and A198P, as well as an insertion of C after A27 and a deletion of I129 (-ATC) (Tables 1 and 2). The overall frequency distribution of unique mutations was as follows: *fbiC* (55%, *n* = 54), *fbiA* (15%, n = 15), *ddn* (12%, n = 12), *Rv2983* (10%, n=10), *fgd1* (4%, *n* = 4), and *fbiB* (4%, n=4) (Fig. 2 and Table S4). There were no clear associations between pretomanid dose and the mutated gene. Mutations in *fbiC* comprised a higher proportion of those selected in our *in vivo* study compared to the proportion selected in a previous in *vitro* study (26%, *p* = 0.0001)(*22*). On the other hand, mutations in *ddn* (29%) were more frequent after *in vitro* selection than in our mouse models (12%) (p =0.001). *In vitro* mutation frequencies for *fbiA, fgd1* and *fbiB* (19%, 7% and 2%, respectively) were similar to our findings in mice.

**Table 1.** Mutations identified in 89 individual colonies and pooled isolates of pretomanid-resistant *M. tuberculosis* selected in BALB/c mice

| Pretomanid treatment (mg/kg) | Mouse ID | No. of single isolates (n=89) | Gene name, amino acid (aa) length and no. of unique mutations selected (n=54) |  |  |  |  |  |
| --- | --- | --- | --- | --- | --- | --- | --- | --- |
|  |  |  | *Rv3261(fbiA)*<br>331aa (n=7) | *Rv3262(fbiB)*<br>448aa (n=3) | *Rv1173(fbiC)*<br>856aa (n=29) | *Rv3547(ddn)*<br>151aa (n=5) | *Rv0407(fgd1)*<br>336aa (n=2) | *Rv2983(cofC)*<br>214aa (n=8) |
| 0 | M-1 | 2 |  |  |  | -G in aa 38 | K9N |  |
|  | M-2 | 4 |  |  | R133P (2); -A in aa 503 |  | K9N |  |
|  | M-4 | 1 |  |  | W690* |  |  |  |
|  | M-5 | 4 |  |  | +A in aa 426 (4) |  |  |  |
| 10 | M-1 | 8 | A212V |  | L702R (4); C562W | -GC in aa 39 |  | +C in aa 27 |
|  | M-2 | 2 |  |  | 9 bp del (aa 307-309);<br>-TCCCCGATG del in aa 306-308 |  |  |  |
|  | M-3 | 4 |  | L15P | L772W; -C in aa 252 (2) |  |  |  |
|  | M-4 | 5 |  |  |  |  |  | G147C (5) |
|  | M-5 | 2 |  |  | A632V |  |  | R25G |
| 30 | M-1 | 1 |  |  | S169P |  |  |  |
|  | M-2 | 3 | L211P |  |  | W20* |  | A132V |
|  | M-3 | 2 |  |  | R278P (2) |  |  |  |
|  | M-4 | 1 |  |  | W155* |  |  |  |
|  | M-5 | 3 |  |  | Q400*; +T in aa 188 (2) |  |  |  |
| 100 | M-1 | 1 |  |  |  |  |  | -ATC in aa 129 |
|  | M-2 | 3 |  |  | K682E; Y170*; L187R |  |  |  |
|  | M-3 | 2 |  | W397R; L173P |  |  |  |  |
|  | M-4 | 2 |  |  | F744S | -C in aa 93 |  |  |
|  | M-5 | 4 | S219G |  | M776T (3) |  |  |  |
| 300 | M-1 | 7 |  |  |  |  |  | R25S (7) |
|  | M-2 | 4 | Q27* (3); -G in aa 248 |  |  |  |  |  |
|  | M-3 | 4 |  |  | H190N; Q20*; -GC in aa 846 (2) |  |  |  |
|  | M-4 | 4 |  |  | T707M; T301A; -C in aa 247 | 9345 bp del (Rv3540-Rv3550) |  |  |
|  | M-5 | 7 | W79* (6) |  |  |  |  | A198P |
| 600 | M-1 | 1 |  |  | N336K |  |  |  |
|  | M-2 | 5 |  |  | R500* (5) |  |  |  |
|  | M-3 | 1 | D49G |  |  |  |  |  |
|  | M-4 | 2 |  |  |  |  |  | C152R (2) |

Note: \* : stop codon; + or ins: insertion; - or del: deletion; no. in parenthesis next to a mutation: no. of isolates with the same mutation.

**Table 2.** Mutations identified in 64 individual colonies and pooled isolates of nitrioimidazole-resistant *M. tuberculosis* selected in C3HeB/FeJ mice

| Pretomanid treatment (mg/kg) | Mouse ID | No. of single isolates (n=64) | Gene name, amino acid (aa) length and no. of unique mutations selected (n=45) |  |  |  |  |  |
| --- | --- | --- | --- | --- | --- | --- | --- | --- |
|  |  |  | *Rv3261 (fbiA)*<br>331aa (n=8) | *Rv3262 (fbiB)*<br>448aa (n=1) | *Rv1173 (fbiC)*<br>856aa (n=25) | *Rv3547 (ddn)*<br>151aa (n=7) | *Rv0407 (fgd1)*<br>336aa (n=2) | *Rv2983 (cofC)*<br>214aa (n=2) |
| 10 | M-1 | 4 |  |  | -A in aa 331; G194D; W198* |  |  | A68E |
|  | M-2 | 4 |  | -T in aa 353 | -C in aa 363 |  | +A in aa 214 (2) |  |
|  | M-3 | 2 |  |  | -C in aa 20 | L49P |  |  |
|  | M-4 | 2 |  |  | K684T |  |  | Q114R |
|  | M-5 | 1 |  |  | del 42 bp (aa 680-693) |  |  |  |
| 30 | M-1 | 5 |  |  | -A in aa 225; +G in aa 107 (2) | W27* |  |  |
|  |  |  |  |  | IS6110 in 85bp upstream of *fbiC* |  |  |  |
|  | M-2 | 5 | -GC in aa 140 |  | L377P (2); L377P (2, het, p) |  |  |  |
|  | M-4 | 3 |  |  | G148D; W354R | C149Y |  |  |
|  | M-5 | 3 | G273D (2) |  | 1 kb del (*fbiC*+PE12) |  |  |  |
| 100 | M-1 | 5 |  |  |  | IS6110 ins. in aa 131 | G191D (5) |  |
|  | M-2 | 1 |  |  | del 4 bp in aa 75-76 |  |  |  |
|  | M-3 | 3 | -G in aa 47 (2) |  | L685R |  |  |  |
|  | M-4 | 5 | +C in aa 125 |  | A588P (2); -G in aa 52; R322L |  |  |  |
|  | M-5 | 2 | L308P (2) |  |  |  |  |  |
| 300 | M-1 | 5 |  |  | A591D (2); A827G; +C in aa 328 (2) |  |  |  |
|  | M-2 | 5 | Q120P (3) |  | R322L; -T in aa 627 |  |  |  |
|  | M-3 | 2 | Q120P (2) |  |  |  |  |  |
|  | M-4 | 3 |  |  |  | R112W (2); IS6110 ins. in D108 |  |  |
|  | M-5 | 4 | D286A |  | T273K (2) | -G in aa 39 |  |  |

Note: \* : stop codon; + or ins: insertion; - or del: deletion; no. in parenthesis next to a mutation: no. of isolates with the same mutation.

**Table 3.** Distribution of mutation types and frequencies in genes associated with pretomanid resistance, by mouse strain

| Mouse model | Type of mutation | Gene name and amino acid (aa) length |  |  |  |  |  | No. of mutation | Frequency of mutation |
| --- | --- | --- | --- | --- | --- | --- | --- | --- | --- |
|  |  | *Rv3261 (fbiA)* | *Rv3262 (fbiB)* | *Rv1173 (fbiC)* | *Rv3547 (ddn)* | *Rv0407 (fgd1)* | *Rv2983 (cofC)* |  |  |
|  |  | 331aa | 448aa | 856aa | 151aa | 336aa | 214aa |  |  |
| BALB/c | no. of point mutation | 4 | 3 | 15 | 0 | 2 | 6 | 30 | 56% |
|  | no. of indels (ins+del) | 1 | 0 | 8 | 4 | 0 | 2 | 15 | 28% |
|  | no. of stop codon | 2 | 0 | 6 | 1 | 0 | 0 | 9 | 17% |
|  | no. of total mutation | 7 | 3 | 29 | 5 | 2 | 8 | 54 | 100% |
|  | frequency of mutation | 13% | 6% | 54% | 9% | 4% | 15% | 100% |  |
| C3HeB/FeJ | no. of point mutation | 5 | 0 | 12 | 3 | 1 | 2 | 23 | 51% |
|  | no. of indels (ins+del) | 3 | 1 | 12 | 3 | 1 | 0 | 20 | 44% |
|  | no. of stop codon | 0 | 0 | 1 | 1 | 0 | 0 | 2 | 4% |
|  | no. of total mutation | 8 | 1 | 25 | 7 | 2 | 2 | 45 | 100% |
|  | frequency of mutation | 18% | 2% | 56% | 16% | 4% | 4% | 100% |  |

Note: ins: insertion; del: deletion.

**Fig. 2.**
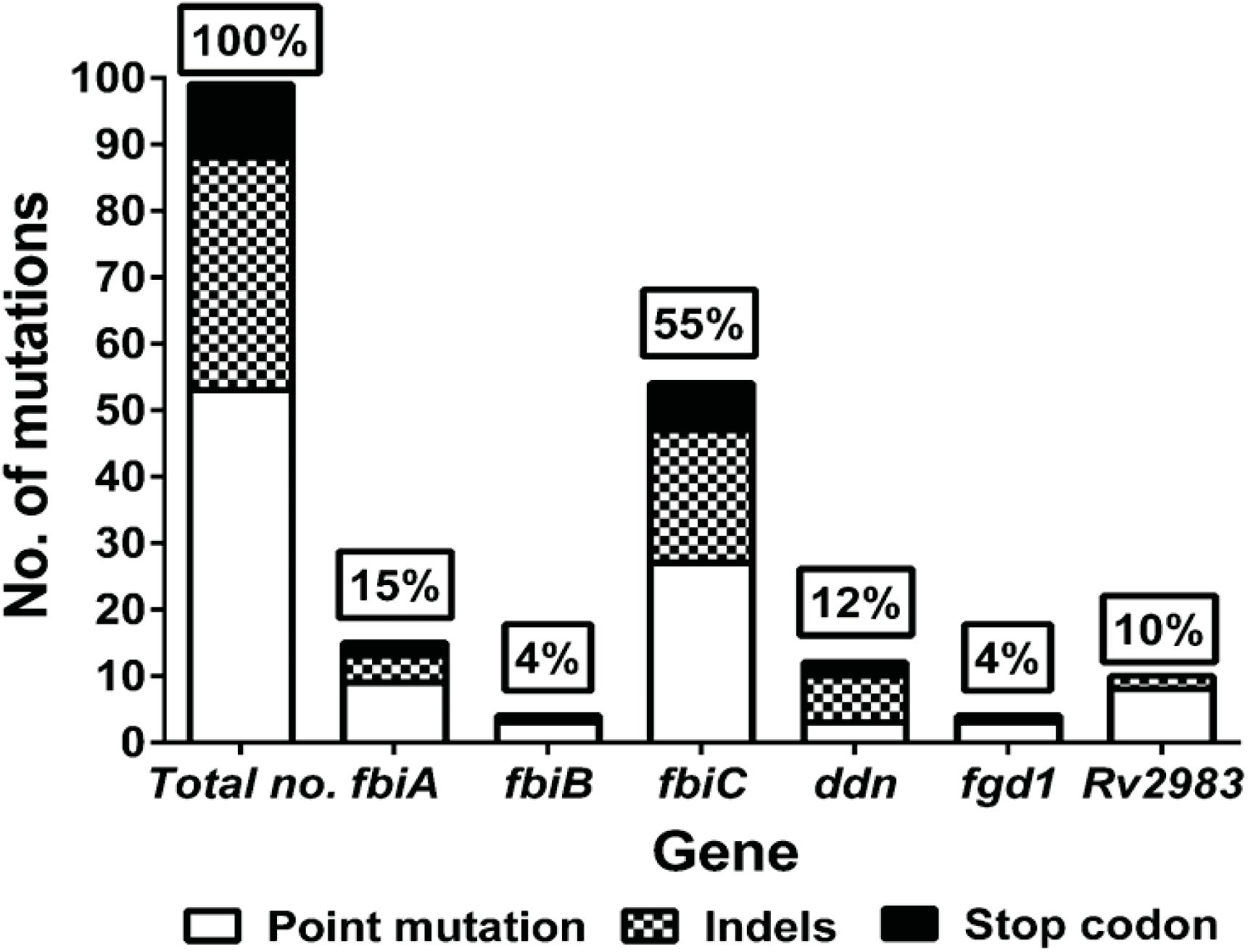
Overall mutation frequencies of genes associated with pretomanid resistance. WGS was performed with 136 pretomanid-resistant colonies and 25 colony pools picked from 47 individual mice harboring pretomanid-resistant CFU after 8 weeks of treatment and identified 99 unique mutations in these 6 genes. Mutations in *fbiC* (55%) were the predominant cause of pretomanid resistance. For the other 5 genes, the rank order by mutation frequency was *fbiA* (15%), *ddn* (12%), *Rv2983* (10%), *fgd1* (4%) and *fbiB* (4%).

Among the 99 unique mutations, all but one (an IS6110 insertion located in 85-bp upstream of the *fbiC* coding sequence in isolate KA-026a (Table 2 and Table S2) were found within the coding regions of the six genes. In total, 54% (53/99) were non-synonymous point mutations (no synonymous point mutations were identified), 35% (35/99) were insertions or deletions (indels), and 11% (11/99) were substitutions resulting in a stop codon. No significant difference in the distribution of point mutation and indels was found between ours and the *in vitro* study by Haver, *et al*, in which non-synonymous point mutations and indels were 50% (75/151) and 24% (36/151), respectively. However, the frequency of stop codon substitutions in the latter study (26%, 40/151) was higher than that observed in the present study (11%, 11/99) (*p* = 0.004), 85% (34/40) of which were in *ddn* in the latter study (22). Non-synonymous point mutations predominated relative to indels and stop codon mutations overall and in each gene except for *ddn* (Fig. 2 and Table S4). The frequency of point mutations was similar between BALB/c and C3HeB/FeJ mice (56% versus 51%) (Table 3). A higher frequency of indels occurred in C3HeB/FeJ compared to BALB/c mice (44% versus 28%), while more stop codon mutations occurred in BALB/c, but these differences were not statistically significant. There was no clear association between dose of pretomanid and the type of mutation selected.

Comparing the 99 unique mutations identified in our study with the 151 unique mutations in 5 of the same genes selected *in vitro* (*22*), only 4 mutations were found in the same position. Only W79 stop (*fbiA*) and N336K (*fbiC*) mutations were found in both datasets while both T273 and H190 (*fbiC*) were mutated in the same position but with different mutations. As expected from the fact that most mutants could be isolated on plates containing 10 μg/ml of pretomanid, MICs determined against a small subset of isolates indicated high-level pretomanid resistance (Table S5 and Table 1 and 2). Taken together, these data illustrate the tremendous diversity of mutations capable of conferring high-level pretomanid resistance.

### Mutations in *Rv2983* cause resistance to pretomanid and delamanid

To prove that mutations in *Rv2983* are sufficient for nitroimidazole resistance, merodiploid complemented strains were constructed by introducing a copy of the wild type *Rv2983* gene into B101, an *Rv2983* mutant (A198P), through site-specific integration (*31, 32*). Following confirmation of successful integration by Southern blot using a DIG-labeled *Rv2983* probe (Figs. S1A and S1B), susceptibility testing by 7H9 broth dilution confirmed significantly higher nitroimidazole MICs against the *Rv2983* mutant (pretomanid and delamanid MICs of 32 μg/ml and 0.064-0.128 μg/ml, respectively) and full restoration of susceptibility in the complemented strains (pretomanid and delamanid MICs of 0.25 μg/ml and 0.008 μg/ml, respectively).

### *Rv2983* is required for F_420_ biosynthesis

To demonstrate that *Rv2983* is required for F_420_ biosynthesis, we measured the production of Fo and F_420_ in *M. smegmatis* strains overexpressing *Rv2983* and in *M. tuberculosis Rv2983* mutant strains compared with their corresponding control strains. *Rv2983* and *fbiC* were successfully cloned into pYUBDuet and *pfbiC* (designated *pRv2983* and *pfbiC-Rv2983*, respectively), followed by successful transformation of *M. smegmatis*, along with pYUBDuet and *pfbiABC*, which were confirmed by restriction enzyme digestion, DNA sequencing and PCR amplification (data not shown). Overexpression of *Rv2983 in M. smegmatis* increased F_420_ production but resulted in little change in Fo production compared to the control strain after 6 and 26 hours of induction with IPTG (Figs. 3A and 3B). As expected, mutation of *Rv2983* in the *M. tuberculosis* B101 mutant markedly reduced F_420_ production, resulting in accumulation of Fo. Complementation fully restored the wild-type phenotype (Figs. 3C and 3D). In order to evaluate the method, we also overexpressed *fbiC*, which encodes the Fo synthase, with and without concomitant overexpression of Rv2983 in *M. smegmatis*. As expected, overexpression of *fbiC* increased Fo and, consequently, F_420_ concentrations. Relative to the control strain, F_420_ concentrations were similar when either *fbiC* or *Rv2983* was over-expressed alone (Fig. 3A). Interestingly, when *Rv2983* was co-overexpressed with *fbiC*, a dramatic increase in F_420_ was observed relative to over-expression of either gene alone (3.4 and 3.1-fold, respectively) after 6 hours of IPTG induction (*p*<0.001), with corresponding significant decreases of Fo levels after 6 and 26 hours of IPTG induction (5.8 and 3.1-fold; *p*<0.005 and 0.05, respectively) (Fig. 3A and 3B). These results suggest that the excess Fo produced by *fbiC* over-expression was efficiently converted to F_420_ by over-expressed Rv2983. On the other hand, although a small amount of F_420_ was observed in cell extracts of two *Rv2983* point mutants (B101 [A198P] and KA016 [Q114R]), their F_420_ content was significantly lower than that of the wild type (7.3 and 7.7-fold) (*p*</italic> < 0.001) and complemented B101 mutant (Fig. 3C). As expected, Fo accumulated in the two *Rv2983* mutant strains relative the wildtype (6.7 and 6.5-fold; *p*<0.05 and 0.005, respectively) (Fig. 3D), indicating that Fo was not efficiently converted to F_420_ in the presence of a mutated *Rv2983*. Two other pretomanid-resistant strains were also assessed as controls. The KA026 mutant with an IS6110 insertion 85 bp upstream of *fbiC* had undetectable Fo and very little F_420_ content, while the KA91 mutant with an IS6110 insertion at amino acid position 108 of Ddn showed a wild-type phenotype with respect to F_420_ and Fo concentrations (Fig.3C and D).

**Fig. 3.**
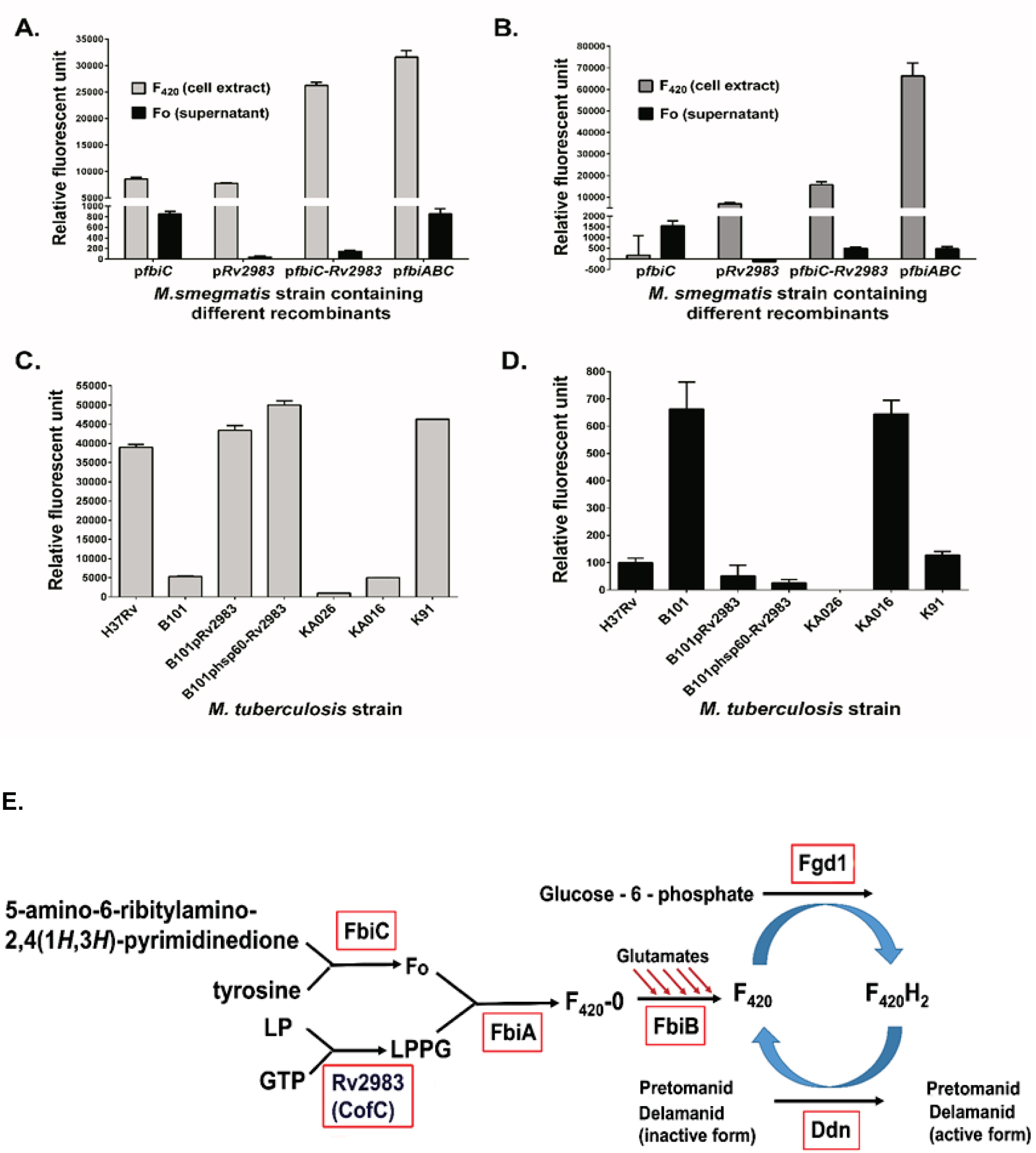
Rv2983 is required for efficient F_420_ synthesis from Fo. F_420_ and Fo content was measured in *M. smegmatis* strains harboring different recombinants relative to the control strain containing the empty vector pYUBDuet after 6 (A) and 26 (B) hours of 1mM IPTG induction; F_420_ (C) and Fo (D) content was measured in *Rv2983* mutant strains of *M. tuberculosis* and control strains including B101 (Δ*Rv2983*, A198P), KA016 (Δ*Rv2983*, Q114R), H37Rv (wild-type), B101 complemented strain (pMH94-Rv2983), B101 complemented strain (pMH94-hsp60-Rv2983), KA026 (*AfbiC*, IS6110 insertion in 85-bp upstream of *fbiC*), and K91 (Δ*ddn*, IS6110 insertion in aa D108), after growth in 7H9 broth for 6 days. Schematic diagram (E) of proposed nitroimidazole activation pathway showing Rv2983 as putative CofC catalyzing LPPG biosynthesis. Fo, 7,8-didemethyl-8-hydroxy-5-deazariboflavin; LP, 2-phospho-L-lactate; GTP, guanosine triphosphate; LPPG, L-lactyl-2-diphospho-5’-guanosine.

To understand the effect of mutations on gene expression and its regulation, we performed RT-qPCR after sub-culturing the *M. tuberculosis* strains in fresh 7H9 broth. Expression of *Rv2983* and *fbiC* increased 2.4‐ and 1.6-fold, respectively, in the *Rv2983* mutant B101 relative to the wildtype H37Rv parent after 4 days of incubation in 7H9 broth. Similar increases were observed in other genes such as *fbiA* (2.3-fold), *fbiB* (2.0-fold) and *fgd1* (2.2-fold) involved in F_420_ biosynthesis, suggesting that the reduced F_420_ content caused by the *Rv2983* A198P mutation resulted in upregulation of the F_420_ biosynthesis pathway (Fig. S2A). Expression of *fbiC* in the KA026 mutant decreased 114-fold compared to that in the wild-type H37Rv parent after 2 days of incubation in 7H9 broth, likely resulting from interrupted *fbiC* transcription as a result of the insertion IS6110 at 85-bp upstream and explaining the low levels of both Fo and F_420_ in that mutant (Fig. S2B). A faint band representing the *fbiC* DNA fragment of 937-bp from the *fbiC* KA026 mutant relative to H37Rv further supports this conclusion (Fig. S2C).

### F_420_-deficient pretomanid-resistant mutants are attenuated for growth in the presence of malachite green

Previous work using *M. smegmatis* showed that mutations in MSMEG_5126 (homolog of *fbiC*) and MSMEG_2392 (which shares 69% homology with *Rv2983*) reduce the ability to decolorize and detoxify MG, indicating that F_420_ is necessary for this process (*20*). To evaluate the role of each gene associated with nitroimidazole activation in the susceptibility to MG, log-phase cultures of 10 selected mutants were plated on 7H9 agar supplemented with a range of MG concentrations. All mutants deficient in F_420_ synthesis or reduction (*i.e.*, those with mutations in *fbiA-C*, *Rv2983* or *fgd1*) were more susceptible to MG, while the *ddn* mutant retained the same susceptibility as the wild type H37Rv parent, whose growth was almost completely inhibited at MG concentrations ≥ 30 μg/ml (Fig. 4A). The lability of F_420_H_2_ and lack of a commercial source for F_420_ made it unfeasible to attempt to test whether provision of F_420_H_2_ could rescue the MG-hypersusceptible phenotype of the F_420_H_2_–deficient mutants. To confirm that *Rv2983* is necessary for the intrinsic resistance of *M. tuberculosis* to MG, we compared the growth of the B101 mutant to that of the wild type and complemented strains on MG. Again, growth of this *Rv2983* mutant was inhibited by lower concentrations of MG than the wild type strain and required longer incubation times before colonies appeared on MG-containing plates (Fig. 4B-D). Plating at higher bacterial density (500 μl rather than 100 μl of cell suspension per plate) significantly increased recovery (Fig. S3A and B). Complementation of *Rv2983* fully restored the wild-type growth phenotype on MG concentrations up to 6 μg/ml. Interestingly, at MG concentrations above 6 μg/ml, greater recovery was observed when *Rv2983* was expressed behind the native promoter compared to the *hsp60* promoter (Fig. 4B-D). Smaller colony size was observed in the *Rv2983* mutant relative to the wild type and the complemented strains on plates without MG after 21 days of incubation but not after 28 days of incubation (Fig. 5). However, smaller colony size and deficient decolorization was observed for the *Rv2983* mutant on plates containing MG concentrations as low as 0.25 and 1 μg/ml, even after 28 days of incubation (Fig. 5). At MG concentrations of 6-12 μg/ml, even more-concentrated aliquots of the B101 mutant culture showed markedly reduced recovery despite 42 days of incubation (Fig. 5).

**Fig. 4.**
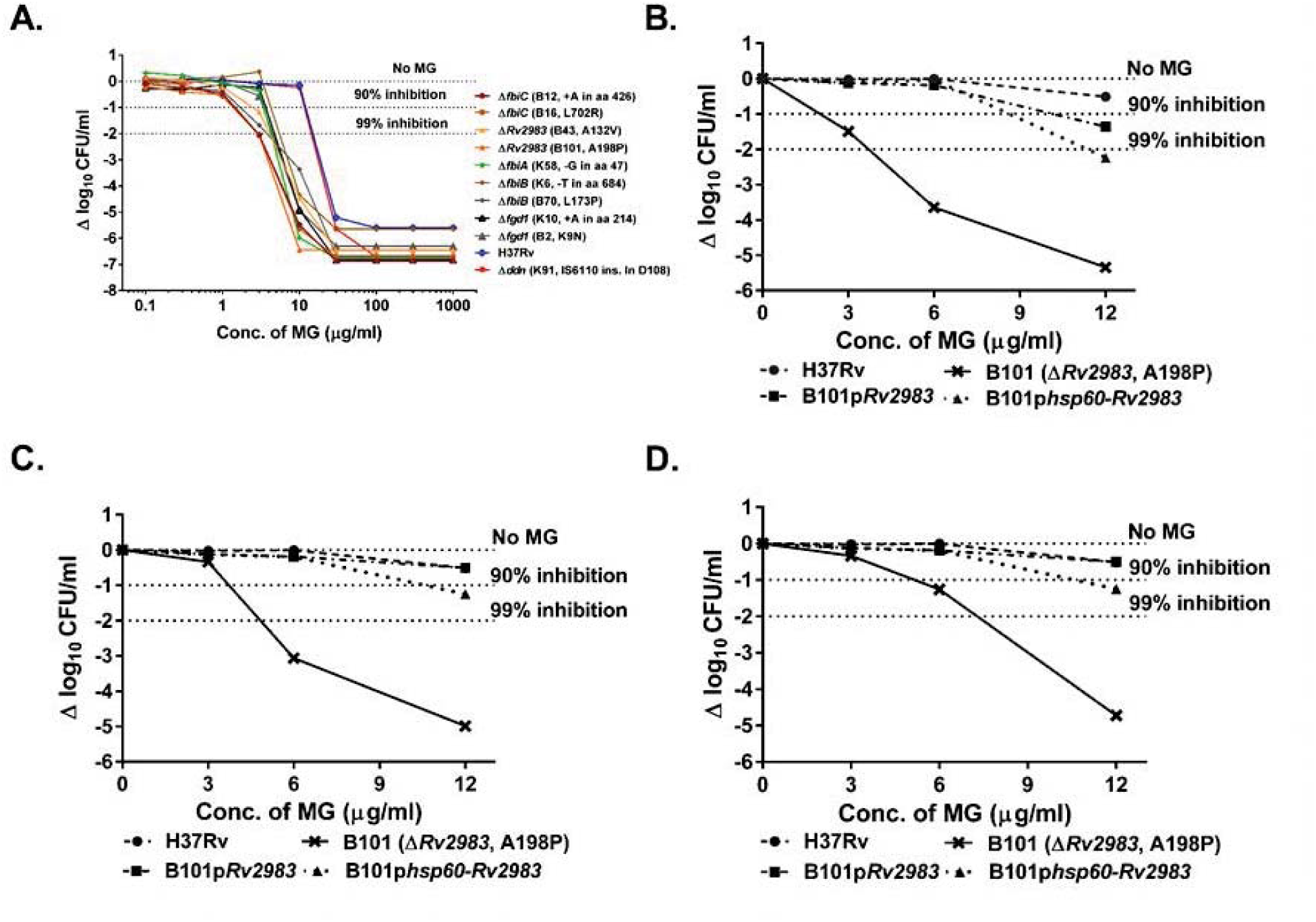
F_420_H_2_-deficient pretomanid-resistant mutants of *M. tuberculosis* are more susceptible to growth inhibition by malachite green. A. Growth of wild-type *M. tuberculosis* on 7H9 agar is inhibited by malachite green (MG) in a concentration-dependent manner. F_420_H_2_–deficient, pretomanid-resistant *M. tuberculosis* mutants (*fbiA-C*, *fgd1*, *Rv2983*) are inhibited at lower MG concentrations relative to the wild type and the F_420_H_2_-sufficient, pretomanid-resistant *ddn* mutant. B-D. Complementation of the B101 mutant with wild-type *Rv2983* restores tolerance to MG and the proportional recovery of the mutant on 6 μg/ml of MG increases as the duration of incubation increases from 28 (B), to 35 (C) to 49 (D) days of incubation. The proportional recovery of the mutant on plates containing 12 μg/ml of MG does not increase with time.

**Fig. 5.**
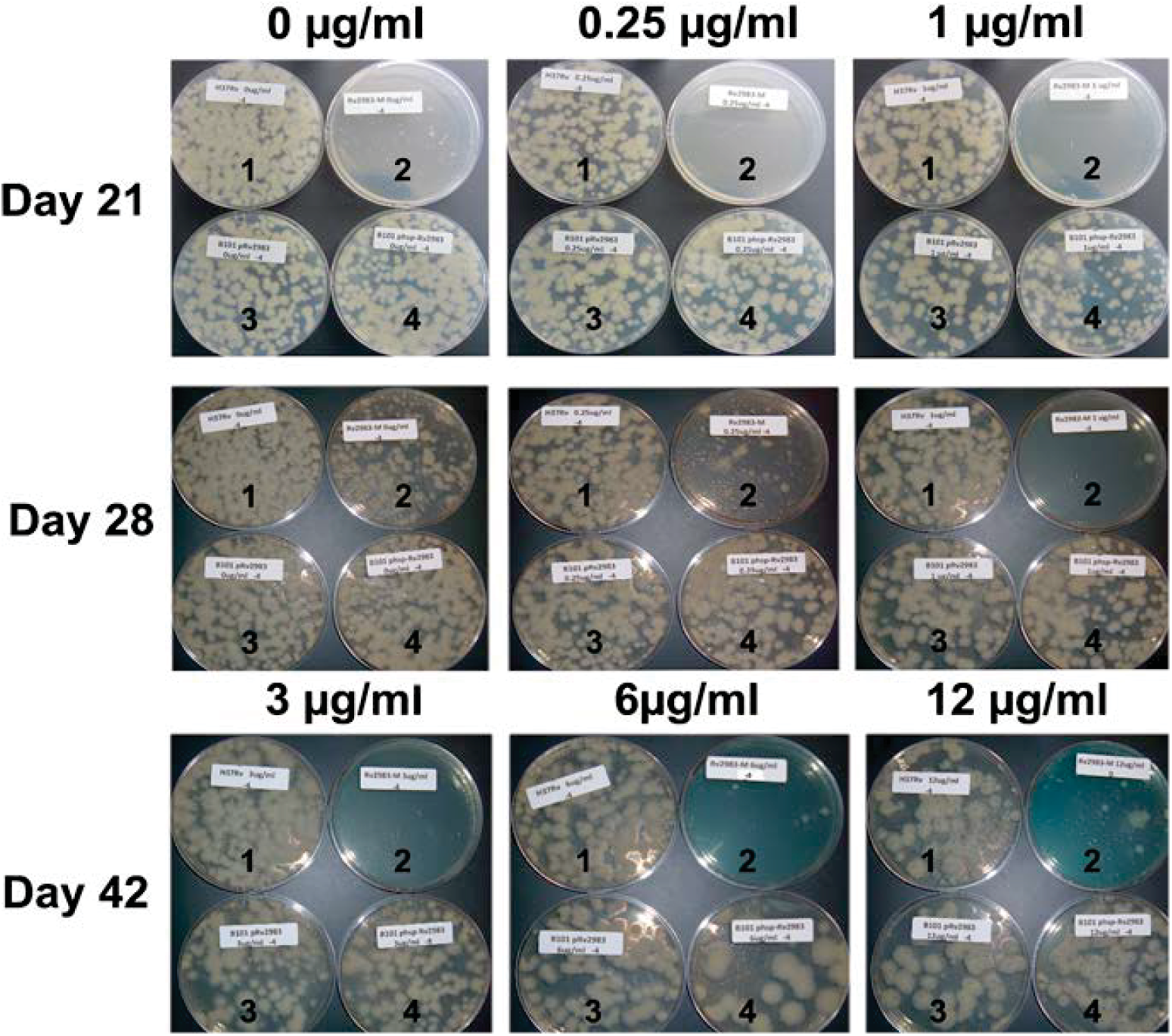
Mutation of *Rv2983* causes growth inhibition and defective decolorization of malachite green. Aliquots (100 μl) of *M. tuberculosis* cultures (1, H37Rv wild type; 2, *Rv2983* mutant B101; 3, B101 mutant complemented with *Rv2983* behind native promoter; 4, B101 mutant complemented with *Rv2983* behind *hsp60* promoter) were spread on 7H9 agar plates containing increasing concentrations of malachite green (MG) after serial 10-fold dilutions. Colony size and MG decolorization are depicted after different incubation times. Numbers on plate labels (e.g., 0, 3, 4, 5) indicate the number of 10-fold dilutions performed before an aliquot was plated. Although the B101 mutant grows more slowly on plates without MG, growth is further slowed in the presence of ≥0.25 μg/ml of MG and the proportion of CFU recovered declines with increasing MG concentrations above 0.25 μg/ml. For plates containing 6 and 12 μg/ml of MG, plates receiving more-concentrated aliquots of the B101 mutant culture are shown to demonstrate the markedly reduced recovery of the mutant at these MG concentrations.

Because all solid media commonly used to isolate and cultivate *M. tuberculosis* in clinical laboratories contain MG as a selective decontaminant, the increased MG susceptibility conferred by mutations in *fbiA-C*, *Rv2983* and *fgd1* could compromise the isolation and propagation (and hence identification) of nitroimidazole-resistant mutants from clinical samples. Commercial 7H10 agar, 7H11 agar and LJ medium contain 0.25, 1 and 400 μg/ml, respectively, of MG. To assess the potential impact of these media on the isolation of an F_420_H_2_-deficient nitroimidazole-resistant *Rv2983* mutant relative to an F_420_H_2_-sufficient, but still nitroimidazole-resistant, *ddn* mutant and the nitroimidazole-susceptible wild type and Rv2983-complemented mutant, we inoculated these media in parallel using serial dilutions of each strain. The *Rv2983* mutant exhibited 10 times lower CFU counts and smaller colony size relative to other strains after 21 and 28 days of incubation on 7H10 agar plates (*p* <0.01) (Figs. 6A and C(a)). The result after 35 days of incubation was generally similar between the mutant and the control strains (Fig. 6A and C(b)). A similar semiquantitative growth assessment of the *Rv2983* mutant on LJ media compared to other strains including a *ddn* mutant (K91, IS6110 ins in D108) revealed growth inhibition of the *Rv2983* mutant that was ameliorated by increasing the size of the bacterial inoculum from 10^2^ to 10^6^ CFU/ml and increasing the incubation time from 28 to 35 days (Fig. 6D). Interestingly, no difference in growth was found on 7H11 agar (Fig. 6B), even when comparing colony size after just 21 days of incubation (Fig. 6C(c)), despite higher MG concentrations in that medium compared to 7H10. Unlike 7H10, 7H11 medium contains hydrolysate of casein, which may somehow mitigate against the MG toxicity. Taken together, these results indicate that use of 7H10 and LJ could compromise the recovery and propagation of F_420_H_2_-deficient nitroimidazole-resistant mutants from clinical specimens and that 7H11 agar may be the preferred solid media for processing specimens propagating any isolates from nitroimidazole-treated patients.

**Fig. 6.**
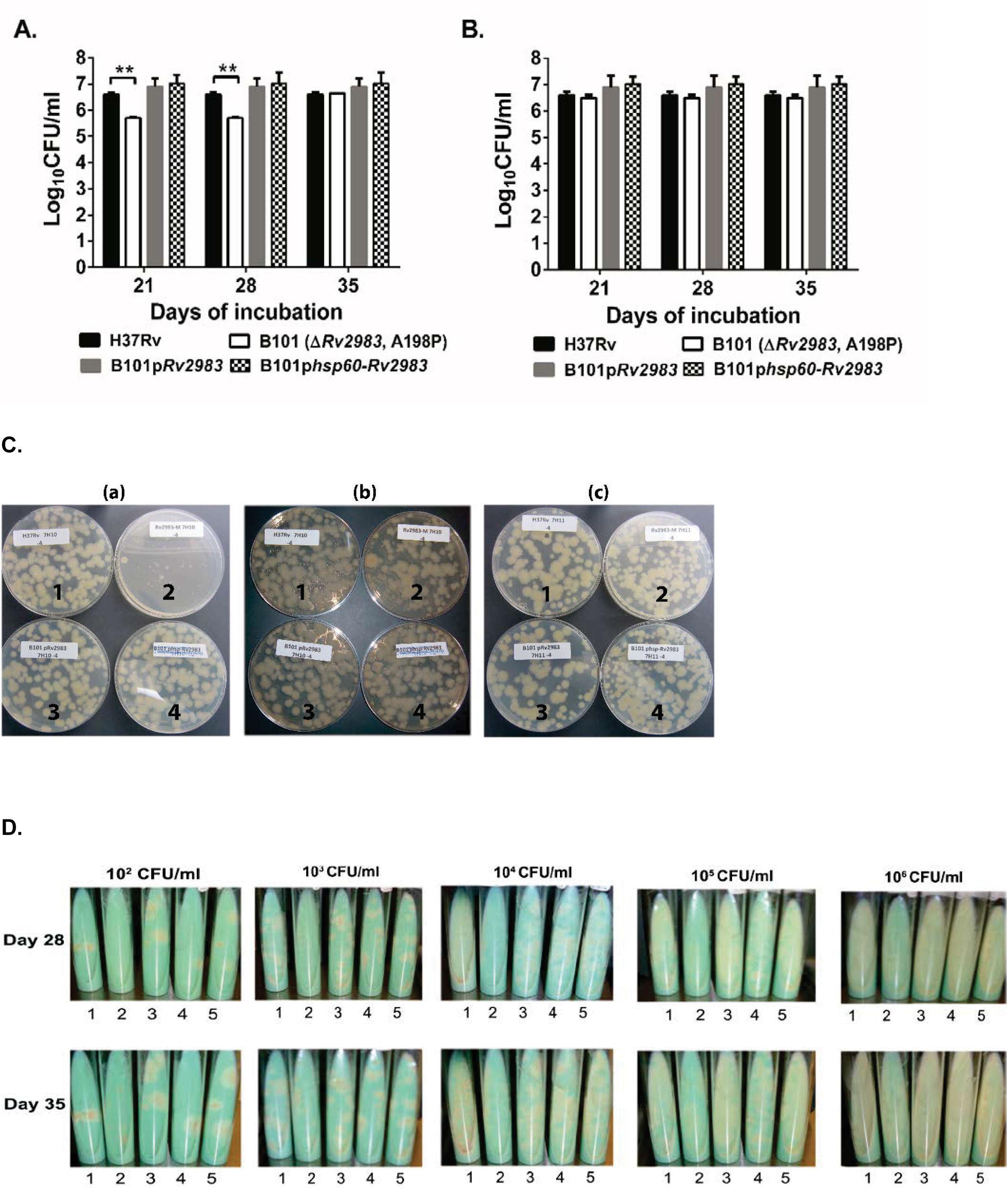
Mutation of *Rv2983* causes growth inhibition on commercial 7H10 agar and LJ slants, but not on commercial 7H11 agar. Aliquots of *M. tuberculosis* cultures were spread on various solid media purchased commercially after serial 10-fold dilutions. A-B. Mean CFU counts on 7H10 (A) and 7H11 (B) agar plates; C. Colonies on 7H10 agar plates after 21 (a) and 35 (b) days of incubation, and on 7H11 agar plates after 21 (c) days of incubation; D. Colonies on LJ slants inoculated with serially diluted aliquots after 28 and 35 days of incubation. 1: H37Rv wild type; 2: B101 mutant (Δ*Rv2983*, A198P); 3: B101 mutant complemented with *Rv2983* behind the native promoter; 4: B101 mutant complemented with *Rv2983* behind the *hsp60* promoter; 5. K91 mutant (Δ*ddn*, IS6110 ins in D108).

## Discussion

As representatives of one of only two new drug classes approved for use against TB in roughly 50 years, delamanid and pretomanid are important and promising new drugs (*3, 4, 6, 7, 39*). The former received accelerated approval from the EMA for treatment of MDR-TB and is now used clinically, albeit sparingly, while a phase 3 trial is being completed. The latter is being evaluated in clinical trials as a component of highly promising regimens for both drug-susceptible and MDR/XDR-TB. Comprehensive knowledge of genetic mutations conferring nitroimidazole resistance in *M. tuberculosis* and the resultant mutant phenotypes are critical for timely and accurate diagnosis of resistance and, therefore, the safe and effective use of these drugs in the clinical setting. Previous work identified 5 genes (*fbiA-C*, *fgd1*, and *ddn*) involved in the activation pathway of nitroimidazole prodrugs in which mutations may confer drug resistance in *M. tuberculosis* complex (*16, 18, 21-23, 40*). In a prior study of the spectrum of nitroimidazole resistance-conferring mutations, Haver, *et al* found that 151 (83%) of 183 pretomanid-resistant isolates selected *in vitro* harbored a single mutation in one of these 5 genes (*22*). However, 17% of the selected strains harbored no mutations in these genes. The present study has several important new findings. First, we identified a novel nitroimidazole resistance determinant—mutations in *Rv2983*—that explained, in the case of our study, all of the pretomanid resistance that was not attributable to mutations in the 5 previously described genes. Second, we demonstrate for the first time that *Rv2983* is required for F_420_ biosynthesis and likely plays a role similar to *cofC* in the methanogen *M. jannaschii* (*19*). Third, we show that *Rv2983* and the ability to produce F_420_H_2_ are essential for full tolerance of *M. tuberculosis* to the selective decontaminant MG, which raises serious concerns about the ability to reliably recover most nitroimidazole-resistant mutants from clinical samples on the most widely used microbiology media.

Our study provides the first comprehensive analysis of the spectrum of nitroimidazole-resistant mutants selected *in vivo* and, because we used WGS, it represents the most comprehensive analysis made to-date. Despite the insignificant differences in mutation frequencies between the two mouse strains for other genes related to nitroimidazole activation, *Rv2983* mutations were observed with a somewhat higher frequency in BALB/c mice, indicating a possible host-specific fitness cost that warrants further dedicated study. Like the *in vitro* study by Haver *et al* (*22*), we found that isolated mutations in *fbiA-C*, *fgd1*, or *ddn* explained the majority of the pretomanid-resistant isolates we selected. However, whereas their study left 17% of resistant isolates unexplained, we found that all of the remaining resistant isolates, representing 10% of the total number of unique mutations, harbored mutations in *Rv2983*, a gene not previously implicated in nitroimidazole resistance. Indeed, the proportion of resistant isolates explained by *Rv2983* (10%) was similar to the proportion explained by *fbiA* (15%) and *ddn* (12%) mutations, which lagged only mutations in *fbiC* (55%) as the predominant cause of pretomanid resistance in our mice. Although the *Rv2983* A198T mutation caused a smaller upward shift in the delamanid MIC compared to the pretomanid MIC, the delamanid MIC of 0.064-0.128 μg/ml against the mutant was still significantly higher than the recently proposed critical concentration of 0.016 μg/ml (*41*). Thus, the identification of *Rv2983* mutations should be included in rapid molecular drug susceptibility tests and algorithms for the diagnosis of nitroimidazole resistance from genome sequence data. The 10 mutations in *Rv2983* identified in this study (Table 1 and 2) represent the first step in the process of identifying specific resistance-conferring mutations to inform test development.

Rv2983 has 22% amino acid sequence identity to CofC (MJ0887) of *M. jannaschii*, a putative guanylyltransferase involved in the biosynthesis of coenzyme F_420_ from the precursor Fo. CofC is believed to catalyze the condensation of 2-phospho-L-lactate (LP) and GTP to form lactyl-2-diphospho-5’-guanosine (LPPG) and PP_i_ (*19*). Subsequent condensation of LPPG with Fo catalyzed by CofD (FbiA) forms F_420_–0 and is followed by sequential addition of glutamate residues by CofE (FbiB) to produce F_420_with a poly-glutamate tail (*19*). Up to now, a CofC homolog has not been identified in *M. tuberculosis*. It is known that Fo biosynthesis is catalyzed by FbiC, but it is not clear how Fo is modified by FbiA to form F_420_–0. Bashiri, *et al* tried to understand the mechanism by overexpressing *fbiC* in the saprophyte *M. smegmatis*, which did not dramatically increase F_420_ biosynthesis, suggesting that an additional intermediate is required to form F_420_–0 from Fo (*35*). Using overexpression of *Rv2983* in *M. smegmatis* and *M. tuberculosis Rv2983* mutants, we provide initial evidence that Rv2983 catalyzes an important step required for synthesis of F_420_ from Fo in the pathogen *M. tuberculosis*, which adds to previous evidence that its ortholog MSMEG_2392 is involved in F_420_ biosynthesis in *M. smegmatis* (*20*). The validity of the method used in this study for detection of F_420_ and Fo was demonstrated by showing the expected results with two pretomanid-resistant strains, KA016 and KA026, harboring mutations in *fbiC* and *ddn*, respectively. Although the role of *Rv2983* remains to be confirmed, our results confirm that expression of *Rv2983* is necessary for efficient conversion of Fo to F_420_ (Fig. 3G).

Our results confirm and significantly extend prior *in vitro* work demonstrating the remarkable diversity of mutations capable of conferring high-level nitroimidazole resistance. Among the 99 unique mutations we identified in 47 mice, only 3 mutations (K9N in *fgd1*, R322L in *fbiC* and Q120P in *fbiA*) were found in more than one mouse. Furthermore, by comparing the 99 unique mutations observed in our mice with the 151 unique mutations selected *in vitro* (*22*), the same mutation occurred only twice. Thus, each of the 6 genes now implicated in nitroimidazole resistance appears to be devoid of “hot spots” for such mutations. The frequency of spontaneous mutations conferring nitroimidazole resistance in *M. tuberculosis* has been investigated *in vitro* by several studies and found to range from 1 in 10^5^ to 7 in 10^7^CFU (*21-23, 27, 40, 42*), which is consistent with our findings in the lungs of untreated BALB/c mice. The large “target size” for mutations in 6 non-essential genes drives these high frequencies of spontaneous nitroimidazole-resistant mutants in *M. tuberculosis* populations, which is as high or higher than that for isoniazid and other TB drugs in clinical use. Our unpublished observations suggest that similar frequencies of nitroimidazole-resistant mutants exist in sputum isolates collected from treatment-naive, drug-susceptible TB patients. Delamanid-resistant *M. tuberculosis* has been recovered from patients both before and after delamanid treatment (*10, 43-45*). To date, emergence of resistance has not been described during use of pretomanid in clinical trials, but such use has been restricted to relatively short treatment durations and/or use of highly active companion drugs. Pretomanid resistance has emerged during combination therapy in mouse models (*3, 46*). Thus, the relatively high frequency of spontaneous mutations conferring nitroimidazole resistance and available pre-clinical and clinical data underscore the importance of making validated drug susceptibility testing for this class widely available as clinical usage expands. The unprecedented number and diversity of resistance-conferring mutations demonstrated for nitroimidazole drugs here and by Haver *et al* (*22*), clearly challenges the development and interpretation of rapid molecular susceptibility tests, especially considering that polymorphisms in nitroimidazole resistance genes that represent phylogenetic markers but do not confer pretomanid resistance are well-described (*47, 48*). A similar situation exists for *pncA* mutations and pyrazinamide (PZA) resistance, where an efficient, yet comprehensive method based on saturating mutagenesis for distinguishing single nucleotide polymorphisms conferring resistance was recently described (*49*). A similar analysis of substitutions in the 6 genes related to nitroimidazole resistance would similarly advance the development of drug susceptibility testing using genome sequencing technology.

Amino acid substitutions in the PZA-activating PncA protein that confer PZA resistance are associated with diminished enzymatic activity or reduced abundance of the protein (*49*). Similar mechanisms could explain the overall mechanisms of the 99 mutations we identified in the six genes conferring nitroimidazole resistance. The *Rv2983* A198P or Q114R substitutions may change the conformation of Rv2983 protein structure and reduce its enzymatic activity given that it has no effect on *Rv2983* gene transcription. F_420_ biosynthesis is dramatically decreased while expression of *Rv2983* and other genes increases, perhaps indicating an unknown regulatory mechanism triggered by reduced F_420_ biosynthesis. Mutations identified in *Rv2983* in this study will provide useful information for future studies on structure-function associations of the potential *M. tuberculosis* CofC homolog. The KA026 mutant harboring an IS6110 insertion 85 bp upstream of *fbiC* exhibited significantly decreased expression of *fbiC*, possibly due to interruption of the promoter region, and, consequently, almost complete loss of Fo synthase activity. Hence, the upstream regulatory regions of the six genes also should be considered in future identification of resistance-conferring mutations, although their prevalence in this study was only 1%.

Our findings regarding the heightened susceptibility of F_420_H_2_-deficient mutants to MG pose a previously unappreciated challenge to the development and use of phenotypic testing methods. Indeed, we observed reduced or delayed recovery of a nitroimidazole-resistant *Rv2983* mutant on commercial 7H10 and LJ media that include MG as a selective decontaminant (*50*). Since *fbiA-C* and *fgd1* mutants, as well as a second *Rv2983* mutant, exhibited similar hypersusceptibility to MG in 7H9 agar supplemented with MG, their recovery on 7H10 and LJ is also likely to be affected. Selective growth inhibition of nitroimidazole-resistant strains on media that are commonly used in clinical microbiology laboratories around the world raises serious concern that their recovery from clinical specimens may be impaired to such an extent that it reduces the sensitivity and accuracy of phenotypic drug susceptibility testing, especially for isolates comprised of mixed wild-type and resistant populations. This concern is further amplified by the common practice of performing susceptibility testing (including molecular testing), not on primary samples but, on isolates that have been sub-cultured one or more times on solid media. Such practices may drastically reduce the proportion of (or eradicate) F_420_H_2_-deficient mutants present in the original sample. In addition, efforts to develop MG decolorization assays for detection of drug-resistant TB are expected to be fruitless for these mutants (*51-54*). We did not determine the basis for the greater recovery of F_420_H_2_-deficient mutants on 7H11 vs. 7H10 media despite 4x higher total MG concentrations in the former. The principal differences between these media are the presence of pancreatic digest of casein in 7H11 and lower concentrations of magnesium sulfate countered by the addition of copper sulfate, zinc sulfate and calcium chloride in 7H10. Although this issue clearly requires further study, we presently believe that 7H10 and LJ should not be employed for phenotypic nitroimidazole susceptibility testing and that primary isolation or subculture of any isolate on such media prior to either phenotypic or genotypic susceptibility testing should be avoided whenever possible. When it cannot be avoided, larger inoculum sizes and longer incubation times may increase recovery on 7H10 and LJ. Based on our study, 7H11 agar appears to be the preferred solid medium for recovery of F_420_H_2_-deficient nitroimidazole-resistant *M. tuberculosis*.

A role for cofactor F_420_H_2_ in tolerance of was previously demonstrated (*20*). Whereas mycobacteria are normally capable of decolorizing and detoxifying MG, mutations in the saprophyte *M. smegmatis* orthologs of *fgd1*, *fbiC* and *Rv2983* disrupt this ability and/or reduce the MIC of MG (*20, 55*). We now extend these observations to the pathogen *M. tuberculosis* and also implicate *fbiA* and *fbiB*, but not *ddn*, in MG tolerance. In *Citrobacter*species, an NADH-dependent triphenylmethane reductase catalyzes reduction of MG to colorless leucoMG that lacks antimicrobial activity and is sequestered in the lipid fraction of the cells (*56, 57*). Triphenylmethane reductase has not been identified in mycobacteria. However, the enhanced MG susceptibility of the F_420_H_2_-deficient mutants, but not *ddn* mutants, suggests that one or more analogous, yet unidentified, F_420_H_2_-dependent reductases is responsible for decolorizing and detoxifying MG in mycobacteria.

Prior studies indicated other fitness costs associated with F_420_H_2_ deficiency, such as hypersusceptibility to oxidative and nitrosative stresses (*12, 58, 59*). Nevertheless, we observed selective amplification of F_420_H_2_-deficient mutants in mice over a range of pretomanid doses that included doses producing much higher drug exposures than those produced in patients. Amplification was especially pronounced at higher drug doses, which eliminated the majority of nitroimidazole-susceptible population more effectively, and in C3HeB/FeJ mice. The lack of marked *in vivo* fitness defects is also supported by the fact that the proportion of all unique resistance mutations explained by each mutation did not differ much between the *in vitro* and *in vivo* conditions, except that a higher proportion of *fbiC* mutations predominated in mice, somewhat at the expense of *ddn* mutations (*22*). Likewise, indels and stop codon mutations comprised nearly 50% of the mutations responsible for resistance. This proportion may be biased due to the high drug doses tested in some mice because such mutations are more likely to result in complete loss-of-function and therefore more complete resistance. However, the frequency of such mutations did not appear to change in a dose-dependent manner. Interestingly, most *Rv2983* mutants were selected in BALB/c rather than C3HeB/FeJ mice despite similar mutation rates in other genes between the two mouse strains. Whether this represents a real or a random difference will require further study. Clearly, clinicians must rely on effective combination drug therapy to prevent the selective amplification of nitroimidazole-resistant mutants. Heightened susceptibility to agents causing oxidative stress has been demonstrated for F_420_H_2_-deficient *M. tuberculosis* and *M. smegmatis* mutants (*58*). A more complete understanding of the unique vulnerabilities of such mutants should aid in the design of effective combination regimens that also optimally restrict selection of nitroimidazole-resistant mutants.

In conclusion, using BALB/c and C3HeB/FeJ mice and WGS, we characterized the pretomanid dose-response relationships for bactericidal effect and suppression of drug-resistant mutants and profiled the genetic spectrum of pretomanid resistance emerging *in vivo*. A novel resistance determinant, Rv2983, was identified as essential for F_420_ biosynthesis and activation of the novel TB pro-drugs delamanid and pretomanid. Furthermore, we provide evidence that F_420_H_2_-deficient, nitroimidazole-resistant *M. tuberculosis* mutants are hypersensitive to MG, raising concern that using MG-containing medium could compromise the isolation and propagation of *M. tuberculosis* from clinical samples and therefore hinder the clinical diagnosis of nitroimidazole resistance. These findings have important implications for both genotypic and phenotypic susceptibility testing to detect nitroimidazole resistance, which will be of increasing importance as wider use of delamanid and, if approved, pretomanid, ensues.

## Acknowledgements

The Global Alliance for TB Drug Development kindly provided pretomanid and delamanid.

## Author contributions

D.R. and E.N. conceived the study and designed the experiments. S.L. and J.L. assisted with the design and conduct of the *in vivo* experiment. Whole genome sequencing was performed and analyzed by T.I., J.S., D.R. and E.N. *In vitro* experiments were performed and analyzed by D.R., J.L. and E.N. The manuscript was drafted by D.R. and E.N. with critical input from T.I., J.S., and G.B.

## Data and materials availability

All data necessary for evaluation of the conclusions are present in the paper and/or the Supplementary Materials. Pretomanid and delamanid were provided under a material transfer agreement.

## Supplementary Materials

**Table S1.**
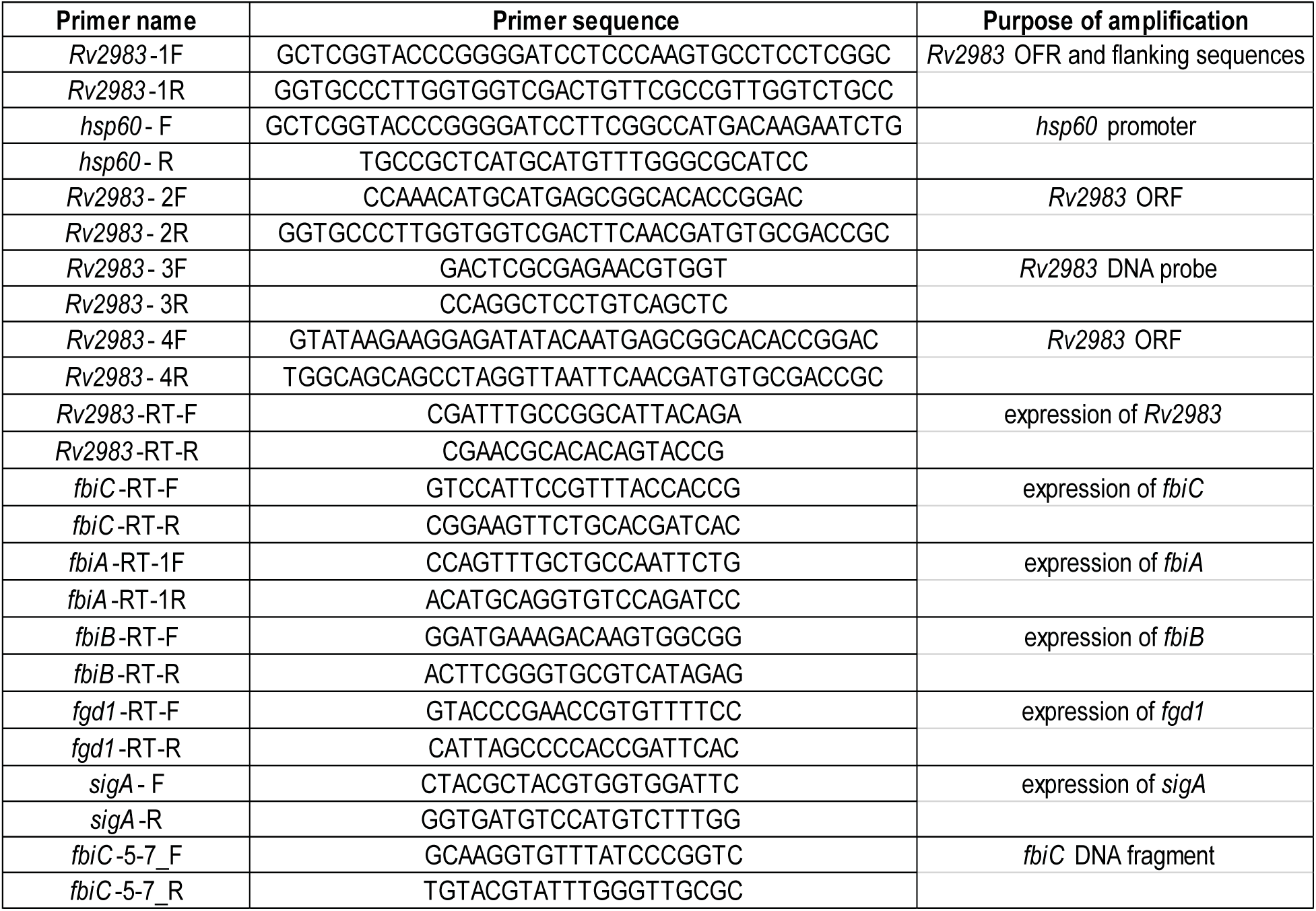
List of the primers used in the study

**Table S2.**
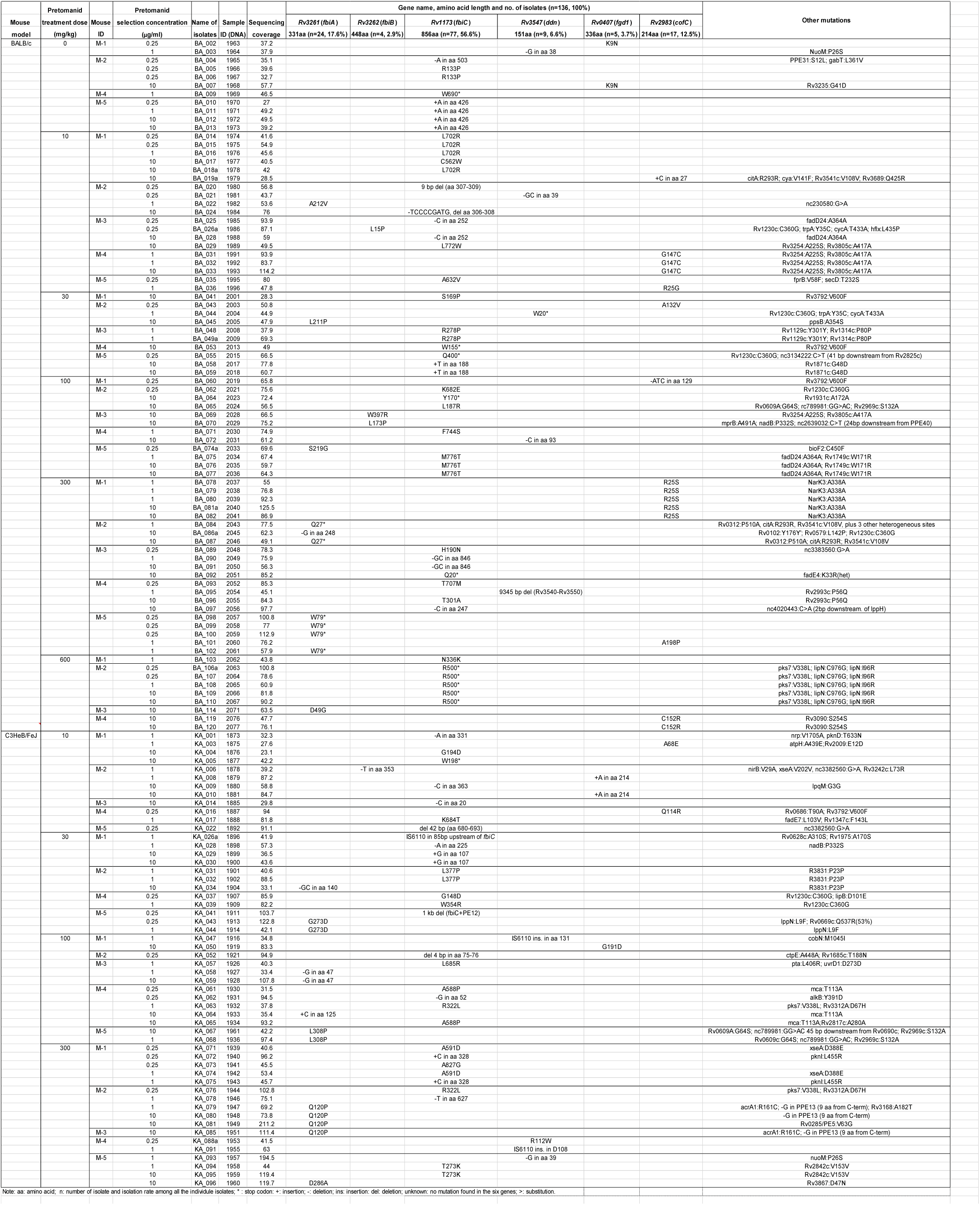
WGS results for 136 individual pretomanid-resistant colonies selected with different doses of pretomanid in *M. tuberculosis-infected* BALB/c and C3HeB/FeJ mice

**Table S3.**
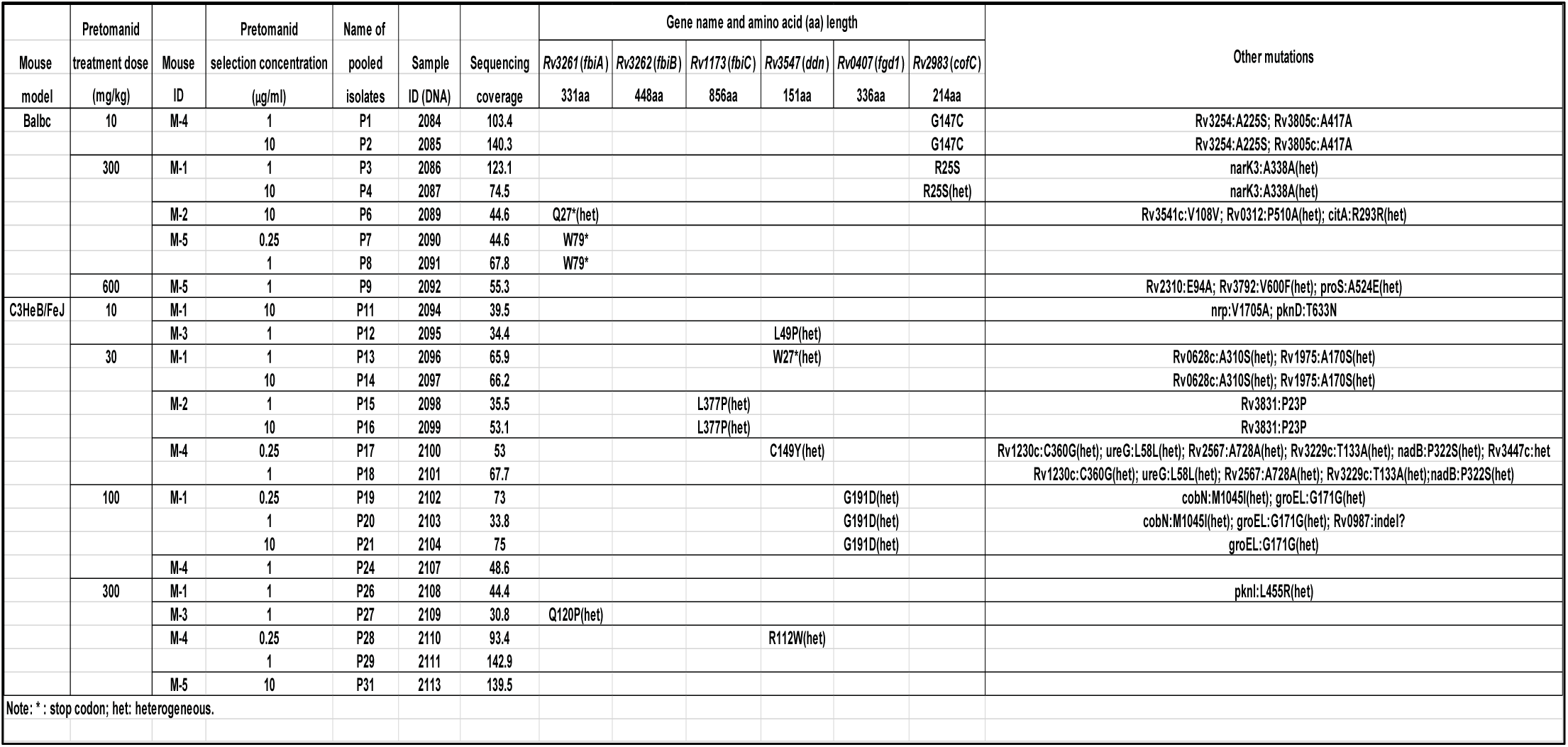
WGS results for 25 pooled pretomanid-resistant isolates selected with different doses of pretomanid in *M. tuberculosis-infected* BALB/c and C3HeB/FeJ mice

**Table S4.** Distribution of overall mutation types and frequencies in genes associated with pretomanid resistance

| Type of mutation | Gene name and amino acid (aa) length |  |  |  |  |  | No. of mutation | Frequency of mutation |
| --- | --- | --- | --- | --- | --- | --- | --- | --- |
|  | Rv3261 (fbiA) | Rv3262 (fbiB) | Rv1173 (fbiC) | Rv3547 (ddn) | Rv0407 (fgd1) | Rv2983 (cofC) |  |  |
|  | 331aa | 448aa | 856aa | 151aa | 336aa | 214aa |  |  |
| no. of point mutation | 9 | 3 | 27 | 3 | 3 | 8 | 53 | 54% |
| no. of indels (ins+del) | 4 | 1 | 20 | 7 | 1 | 2 | 35 | 35% |
| no. of stop codon | 2 | 0 | 7 | 2 | 0 | 0 | 11 | 11% |
| no. of total mutation | 15 | 4 | 54 | 12 | 4 | 10 | 99 | 100% |
| frequency of mutation | 15% | 4% | 55% | 12% | 4% | 10% | 100% |  |

Note: ins: insertion; del: deletion.

**Table S5.** Pretomanid MICs against selected pretomanid-resistant *M. tuberculosis* mutants

| Mouse model | Isolate | Pretomanid treatment dose (mg/kg) | Pretomanid selection concentration (µg/ml) | Gene | Mutation | Agar-based MIC (µg/ml) | Broth-based MIC (µg/ml) |
| --- | --- | --- | --- | --- | --- | --- | --- |
|  | H37Rv parent |  |  |  | wild-type | 0.06 | 0.25 |
| BALB/c | BA_002 | 0 | 0.25 | *fgd1* | K9N | >32 |  |
|  | BA_016 | 10 | 1 | *fbiC* | L702R | >32 |  |
|  | BA_101 | 300 | 1 | *Rv2983* | A198P | 32 |  |
| C3HeB/FeJ | KA_006 | 10 | 1 | *fbiB* | del of T684 | 8 |  |
|  | KA_058 | 100 | 1 | *fbiA* | del of G47 | >32 |  |
|  | KA_091 | 300 | 1 | *ddn* | D108 (IS6110 ins) | 32 |  |
|  | KA-016 | 10 | 0.25 | *Rv2983* | Q114R |  | >32 |
|  | KA-026a | 30 | 1 | *fbiC* | IS6110 ins. in 85bp upstream of *fbiC* |  | >32 |

**Figure S1.**
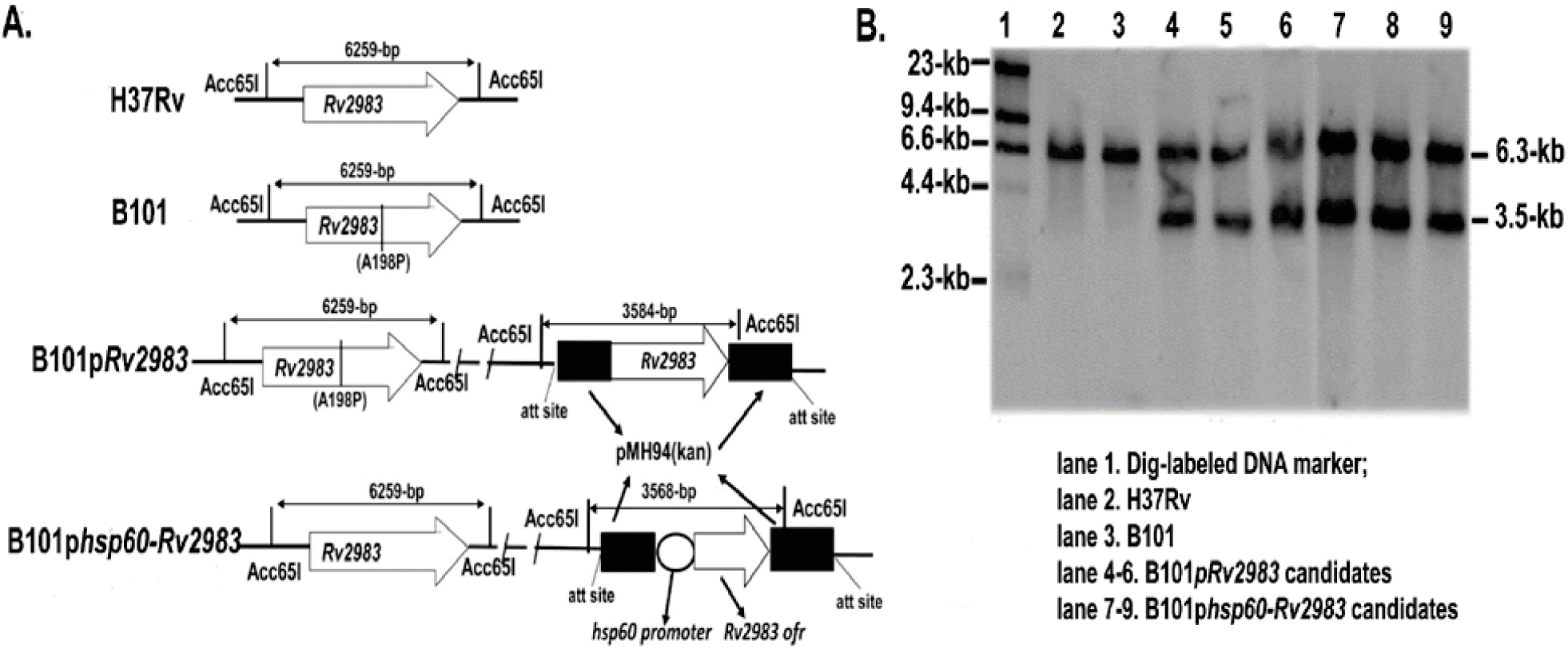
Complementation of B101 mutant with *Rv2983*. A. Schematic diagram of genomic DNA of *M. tuberculosis* strains after digestion with restriction enzyme Acc65I; B. Result of southern blot confirmed expected DNA fragments after Acc65I digestion using DIG-labeled *Rv2983* probe (H37Rv: 6.3 kb; Rv2983 mutant: 6.3 kb; complemented strains: 6.3 and 3.5 kb).

**Figure S2.**
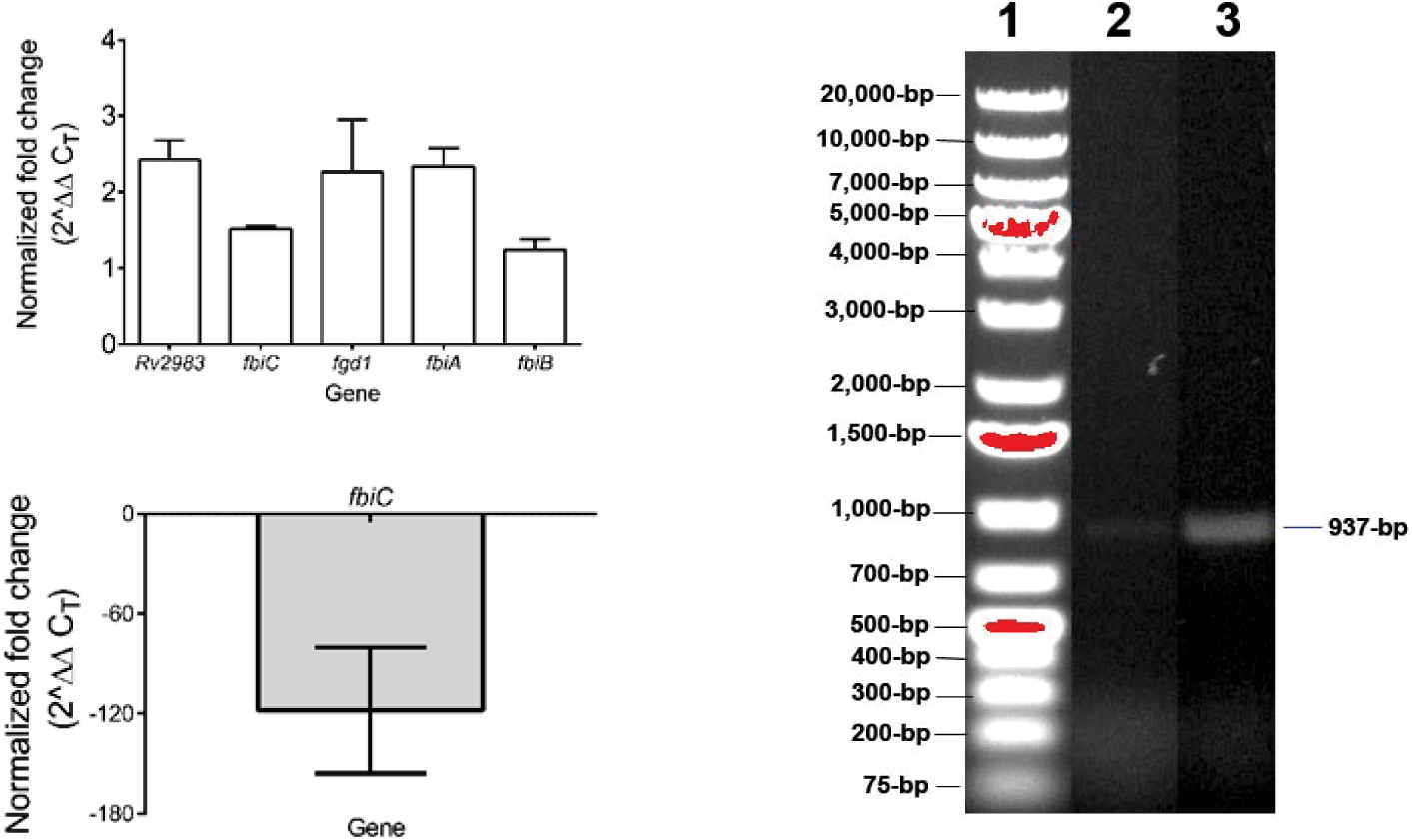
Expression of *Rv2983* and other genes involved in nitroimidazole activation. A. Expression of *Rv2983* and other genes involved in nitroimidazole activation is higher in the *Rv2983* mutant B101 relative to the wild-type H37Rv after 4 days of incubation in 7H9 broth; B. *fbiC* expression is dramatically lower in the *fbiC* mutant KA026 relative to the wild-type after 2 days of incubation in 7H9 broth; C. A faint band representing the 937-bp *fbiC* DNA fragment is evident in the sample from the KA026 mutant (lane 2) relative to that in H37Rv (lane 3). Lane 1 is the 1-kb DNA marker.

**Figure S3.**
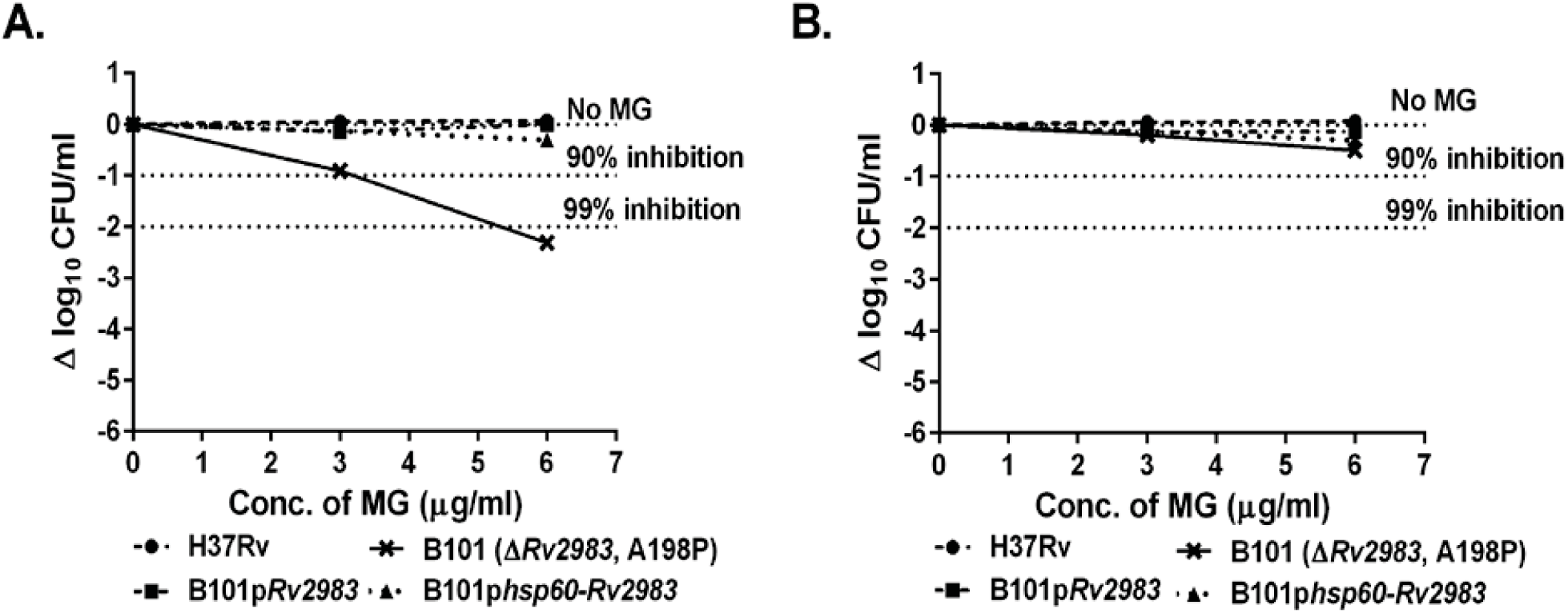
Complementation of the B101 mutant with wild-type *Rv2983* restores tolerance to MG. The proportional recovery of the mutant on 6 μg/ml of MG increases with the volume of culture plated and the duration of incubation: 28-day incubation of 500 μl (A) aliquots/plate; 35-day incubation of 500 (B) aliquots/plate.

